# Deoxyguanosine Kinase Deficiency Couples Purine Metabolism to Innate Immune Activation and Lipid Accumulation in Hepatocytes

**DOI:** 10.1101/2025.11.18.688341

**Authors:** Maija Corey, Mahati Rayadurgam, Jeamin Jung, Alicia Gibbons, Neha Reddy, Mousa Vatanmakanian, Kiyokazu Kakagawa, Priyanka Saminathan, Sonia Sharma

## Abstract

Mitochondrial DNA depletion syndromes (MDS) encompass a heterogeneous set of metabolic disorders caused by defects in enzymes responsible for maintaining mitochondrial nucleotide pools and genome integrity. Among these, deoxyguanosine kinase (DGUOK) acts within the mitochondrial purine-salvage pathway and loss-of-function mutations give rise to DGUOK deficiency, a severe hepatocerebral form of MDS marked by liver failure, neurodevelopmental impairment, and systemic metabolic inflammation. Although the clinical manifestations of DGUOK deficiency have been primarily ascribed to defective mitochondrial DNA (mtDNA) replication, some patients exhibit hepatic steatosis and inflammation despite preserved mtDNA content, suggesting that DGUOK deficiency may deregulate additional metabolic and immune pathways. Here we show that DGUOK depletion reprograms hepatocellular metabolism and innate immune signaling through a purine-dependent mechanism operating independently of mtDNA depletion. In human hepatocellular carcinoma hepatocytes (HEPG2) subjected to siRNA-mediated DGUOK silencing, mitochondrial architecture and respiration remained intact but cells exhibited pronounced lipid droplet accumulation and a robust cell-intrinsic innate immune type I interferon response. Bulk RNA sequencing revealed widespread transcriptional reprogramming, including upregulation of human endogenous retroviruses (HERVs) and interferon-stimulated genes (ISGs), suppression of lipid metabolic pathways, and changes in purine, methionine, and methylation-associated gene networks. Perturbing purine homeostasis through deoxyadenosine (dAdo) supplementation in wild-type cells phenocopied DGUOK disruption, causing global DNA hypomethylation and activation of viral mimicry pathways. Together, these findings identify DGUOK as a central regulator of the purine-regulated lipid-immune axis in hepatocytes, demonstrating that mitochondrial nucleotide salvage preserves hepatic immune and metabolic homeostasis beyond its canonical role in mtDNA synthesis. By linking purine imbalances to steatosis and type I interferon activation, this study establishes a mechanistic framework for immunometabolic pathology in DGUOK deficiency.

## 1 Introduction

Mitochondrial DNA-depletion syndromes (MDS) are a genetically heterogeneous group of autosomal-recessive disorders characterized by a profound reduction in mtDNA copy number and ensuing tissue-specific dysfunction^1^. As mtDNA encodes critical components of the oxidative phosphorylation (OXPHOS) machinery, MDS preferentially impacts tissues with high bioenergetic requirements, such as the liver, skeletal muscle, and central nervous system^1,2^. Disruption of mitochondrial metabolism culminates in hepatic failure, muscle pathology, and neurodevelopmental deficits^1,3^. Liver biopsies from MDS patients commonly display structurally condensed mitochondria and lipid droplet accumulation, reflecting integrated energetic and metabolic stress responses^4,5^.

DGUOK plays a central role in the mitochondrial purine-salvage pathway, catalyzing the phosphorylation of deoxyguanosine (dGuo) and deoxyadenosine (dAdo) to their respective monophosphates, dGMP and dAMP, which are essential precursors for mtDNA synthesis^6,7^. Loss of DGUOK function disrupts mitochondrial dNTP homeostasis and causes a multisystemic hepatocerebral form of MDS that accounts for roughly three-quarters of known DGUOK deficiency cases^8,9^. Clinically, affected infants present with cholestasis, hepatomegaly, and mixed micro- to macrovesicular lipid accumulation (i.e. steatosis) that progresses to liver failure, resulting in death at a mean age of seven months^8,10,11^. Although liver transplantation can restore hepatic function, it offers only partial and transient relief, and approximately half of recipients experience recurrent liver dysfunction and progression of extra-hepatic disease^11,12^.

Emerging clinical and experimental evidence suggests that DGUOK deficiency pathophysiology cannot be explained solely by reduction of mtDNA content. Among 173 reported clinical cases of DGUOK deficiency, a significant reduction in mtDNA copy number was observed in fewer than half of individuals, suggesting that DGUOK contributes to disease pathology through mechanisms beyond mitochondrial genome maintenance^9^. Notably, several DGUOK-deficient patients who did not meet criteria for mtDNA depletion nonetheless developed hepatic steatosis and inflammation^6,10,13^. A remarkable monozygotic twin report highlights a potential interplay between DGUOK loss-of-function and immune dysregulation: despite identical DGUOK mutations, one twin developed Herpes Simplex Virus (HSV)-induced fulminant hepatic necrosis, whereas the uninfected twin displayed subclinical steatosis and mild inflammatory changes^14^. Given that fewer than half of DGUOK-deficient patients exhibit measurable mtDNA copy number reduction yet still develop hepatic injury ^9,13,15^, these observations suggest that DGUOK dysfunction sensitizes hepatocytes to metabolic and immune stress, acting through mechanisms beyond mtDNA depletion.

DGUOK operates within the mitochondrial branch of the purine nucleoside-salvage pathway^6,7,16,17^. In humans, purines and their associated metabolic enzymes are tightly linked to both methyl-donor availability and the activation of innate immune pathways^18–20^. Deficiency of adenosine deaminase enzymes (ADA1 and ADA2), which also catabolize deoxyadenosine (dAdo), perturbs dAdo metabolism which consequently interferes with the cellular methionine cycle, leading to depletion of S-adenosylmethionine (SAM), the universal methyl donor^18^. Reduced SAM availability triggers global DNA hypomethylation and de-repression of human endogenous retrovirus (HERV) elements, whose transcripts engage innate pattern-recognition receptors such as Toll-like receptor 3 (TLR3) and retinoic acid–inducible gene I (RIG-I), culminating in type I interferon (IFN-I) activation and upregulation of interferon-stimulated genes (ISGs)^18,21–23^. Given that DGUOK also functions in purine-salvage metabolism, we reasoned that DGUOK deficiency disturbs both purine balance and methyl-HERV homeostasis, thereby linking mitochondrial metabolism to viral mimicry and antiviral signaling. In hepatocytes, SAM depletion impairs phosphatidylcholine synthesis and very-low-density lipoprotein (VLDL) export, providing a potential mechanistic bridge between purine-driven methyl-donor imbalance and hepatocellular steatosis^24–26^.

Here, we investigated the immediate cellular consequences of DGUOK depletion in hepatocytes, before the onset of mtDNA loss. Using siRNA-mediated silencing of DGUOK in HEPG2 cells, we assessed mitochondrial integrity, transcriptional remodeling, and metabolic reprogramming. Short-term disruption of DGUOK expression triggered type I interferon (IFN-I)-related gene activation and lipid accumulation, while preserving mitochondrial morphology and respiratory function. Bulk RNA sequencing further revealed coordinated enrichment of purine-, SAM-, and methylation-linked pathways, together with upregulation of ISG and HERVs. Importantly, direct perturbation of purine balance via dAdo supplementation in wild-type cells reproduced these effects, inducing global DNA hypomethylation, lipid accumulation, and elevated ISG expression. Collectively, these results identify DGUOK as a previously unrecognized regulator of hepatocellular immune-metabolic homeostasis, extending the framework of MDS beyond genome maintenance to include purine-driven immunometabolic regulation.

## 2 Materials and Methods

### 2.1 Cell Culture

#### 2.1.1 Cell Culture Maintenance

HEPG2 cells (ATCC) were maintained in Eagle’s Minimum Essential Medium (EMEM) (ATCC) with 10% heat inactivated Fetal bovine serum (FBS) and 100 U ml^-1^ penicillin-streptomycin, at 37 °C, 5% CO_2_, and saturated humidity.

#### 2.1.2 siRNA Transfection

siGenome siRNA oligonucleotide pools were purchased from Dharmacon, and transfections were performed using Lipofectamine™ RNAiMAX Transfection Reagent (Thermo Fisher Scientific) according to the manufacturer’s instructions with minor modifications optimized for HepG2 cells. Plates were pre-coated with poly-D-lysine (PDL) to enhance cell adherence and transfection efficiency.

For standard single-gene transfections, 6 pmol of siRNA were used per well of a 24-well plate and 120 pmol per 100 mm dish. In the 24-well format, siRNA–lipid complexes were prepared in 100 µL Opti-MEM™ I Reduced Serum Medium (Thermo Fisher Scientific) with 1 µL Lipofectamine™ RNAiMAX and incubated for 10–20 min at room temperature before addition of 0.05 × 10⁶ cells in 400 µL antibiotic-free complete medium (final siRNA concentration = 10 nM). Reverse transfection was used for initial transfections, as it yielded higher efficiency in HepG2 cells. Cells were harvested at 48 h for RNA analysis, 72 h for protein and functional assays (including lipid quantification), and 96 h for mtDNA quantification, unless otherwise specified.

For experiments involving sequential siRNA transfections, HepG2 cells were first transfected (reverse) with siRNAs targeting innate immune signaling components: IFNAR1 (interferon alpha/beta receptor subunit 1), TMEM173 (STING; stimulator of interferon genes), DDX58 (RIG-I; cytosolic RNA sensor), or RELA (p65; NF-κB subunit). After 48 h, cells were re-transfected (forward) with siRNA targeting DGUOK, either alone or in combination with each innate immune siRNA, to assess pathway-specific modulation of lipid accumulation. A non-targeting siRNA (siCON) served as a negative control. Cells were collected 72 h post-DGUOK transfection for RNA, protein, and lipid imaging analyses.

#### 2.1.3 Plasmid Construct and CRISPR-Cas9 Transfection

The plasmid backbone, Pb-513B-1 (System Biosciences), a PiggyBac vector with a blasticidin resistance cassette and Green fluorescent protein (GFP) reporter gene, was used to construct Single-guide RNA (sgRNA) plasmids targeting DGUOK exons 1, 2, and 3 (sequences: AGGAGGAAACGCCCTCGAGT (-), CTACAGAACCTGTAGCAACA (+), TGTACCGGGAGCCAGCACGA (+)). Plasmids were generated using NEBuilder HiFi DNA Assembly (New England Biolabs) following BbsI digestion, which produced two 5 kb fragments and a ∼22 bp fragment. Fragments were purified using a gel recovery kit(Qiagen), and Gibson assembly was designed to integrate sgRNA sequences into the plasmid and repair the blasticidin cassette cut site. Assembled constructs were transformed into Stbl3 bacteria(Invitrogen), plated on Ampicillin plates, and validated by sequencing.

HEPG2 cells were seeded at 100,000 cells/well in a 24-well plate and transfected with plasmids targeting DGUOK via electroporation (300 ng plasmid DNA/well with 0.076 µL transposase/well). Electroporation was performed using Buffer R (Invitrogen) and settings of 1230-20-3 with the Neon Transfection System (Invitrogen). All three sgRNA constructs were transfected into the same well to maximize targeting efficiency. Post-transfection, cells underwent selection with 1.5 µg/mL treatment of Blasticidin S (Gibco) for six weeks. Knockout (KO) efficiency was assessed by analyzing DGUOK expression at the RNA and protein levels after six weeks.

### 2.2 Total RNA Extraction and RT-qPCR

Total cellular RNA was extracted using Quick-RNA Miniprep Plus Kit (Zymo Research) and cDNA synthesized using qScript cDNA synthesis kit (Quanta), per manufacturer’s instructions. RT–qPCR was performed using CFX96 or CFX384 Touch Detection System (Bio-Rad), with Taqman Universal PCR Master Mix (Applied Biosystems). Messenger RNA abundance of each gene was normalized to ACTB levels in Taqman assay (Thermo Fisher Scientific). Primer sequences are listed in Supplementary Table 1.

### 2.3 Western blotting

For whole-cell lysates, cells were washed in chilled Phosphate-buffered saline (PBS) and lysed in 1× RIPA buffer with Protease/Phosphatase Inhibitor Cocktail (Cell Signaling Technology). After rotation for 30 min at 4 °C, lysates were centrifuged at 15,000 r.p.m. for 15 min to collect supernatants. Protein concentration in supernatants was quantified using Pierce Bicinchoninic acid (BCA) assay Protein Assay Kit, and 50 μg was loaded per well(Thermo Fisher Scientific). Sample preparation, gel electrophoresis, transfer and blocking were performed as previously described^18^. Blots were incubated with DGUOK primary antibody (1:500; Santa Cruz, cat no sc-398101) at 4 °C overnight, with ACTB incubation (1:10,000) at 22 °C for 1 hour. Blots were washed with Tris-buffered Saline-0.1% with Tween 20 and incubated with horseradish peroxidase–conjugated secondary antibodies (1:1000, cell-signaling) at 22 °C for 1 hour. Signal was detected using Enhanced chemiluminescence (ECL) Reagent (Bio-Rad) and imaged on the Bio-Rad Gel Doc.

### 2.4 mtDNA Copy Number Quantification

Total cellular DNA was extracted using the Quick-DNA Miniprep Plus Kit (Zymo Research). Mitochondrial DNA (mtDNA) copy number was quantified by quantitative PCR (qPCR) on a CFX96 or CFX384 Touch Real-Time PCR Detection System (Bio-Rad) using TaqMan Universal PCR Master Mix (Applied Biosystems). MT-CO1 and MT-ND2 were each normalized to the nuclear gene HBB (Thermo Scientific), and relative copy number was calculated using the 2^−ΔΔ^Ct method^27–29^.

### 2.5 Transmission Electron Microscopy

siControl and siDGk transfections were carried out for 72 hours using standard protocols to ensure efficient knockdown of the target gene. HEPG2 WT and KO cells were cultured on 100 mm dishes, with Blasticidin S selection applied for 5 days to ensure the selection of successfully modified cells. Cells were fixed in 2.5% glutaraldehyde prepared in 0.1 M sodium cacodylate buffer and incubated at room temperature for 1 hour, followed by 4 °C incubation for 96 hours. The fixed samples were processed by the University of California San Diego (UCSD) Electron Microscopy (EM) Core Facility. Sample processing involved fixation in glutaraldehyde, incubation in 1% osmium tetroxide, staining with 2% uranyl acetate, dehydration through a graded ethanol series, resin infiltration and polymerization, ultrathin sectioning, and post-staining with 2% uranyl acetate and lead citrate.

Electron microscopy images were analyzed using ImageJ^30,31^ to quantify mitochondrial and lipid droplet morphology^32,33^. Mitochondrial size was defined as the cross-sectional area of individual mitochondria within each cell, while cytoplasmic area was calculated by subtracting the nuclear area from the total cellular area. Lipid droplet accumulation was determined as the total lipid droplet area within the cytoplasm divided by the cytoplasmic area of each cell. Lipid droplet size was defined as the area of individual lipid droplets. Organelle occupancy was expressed as the ratio of organelle area to cytoplasmic area, reported as a percentage.

### 2.6 RNA Bulk-Sequencing Mapping and Differential Expression Analysis

The paired-end reads that passed Illumina filters were filtered for reads aligning to tRNA, rRNA, adapter sequences, and spike-in controls. The reads were then aligned to GRCh38 reference genome and Gencode v27 annotations using STAR (v2.6.1)(PMID: 23104886). DUST scores were calculated with PRINSEQ Lite (v 0.20.3) and low-complexity reads (DUST > 4) were removed from the BAM files^34,35^. The alignment results were parsed via the SAMtools to generate SAM files. Read counts to each genomic feature were obtained with the featureCounts (v 1.6.5) using the default option along with a minimum quality cut off (Phred > 10)^36,37^. After removing absent features (zero counts in all samples), the raw counts were then imported to R/Bioconductor package DESeq2 (v 1.24.0) to identify differentially expressed genes among samples. P-values for differential expression were calculated using the Wald test for differences between the base means of two conditions^38^. These P-values were then adjusted for multiple test corrections using the Benjamini Hochberg algorithm^39^. We considered genes differentially expressed between two groups of samples when the DESeq2 analysis resulted in an adjusted P-value of <0.10 and the difference in gene expression was 1.5-fold. Principal Component Analysis (PCA) was performed using the ‘prcomp’ function in R. The sequencing data generated in this article have been submitted to the Gene Expression Omnibus under accession number GSE309813 (http://www.ncbi.nlm.nih.gov/geo/).

Volcano and MA plots were generated in ggplot2 v3.5.1. Volcano: log2 fold-change (x) vs – log10(adjusted P-value) (y); Gene set enrichment used GSEA-Preranked (classic scoring) with Gene Ontology (GO) gene sets from Molecular Signatures Database (MSigDB); ranks were – log10(P) for positive log2FC and +log10(P) for negative log2FC.

### 2.7 HERV Analysis

For analysis of HERV expression, the paired-end reads that passed Illumina filters were filtered for reads aligning to tRNA, rRNA, adapter sequences, and spike-in controls. The reads were then aligned to the reference genome using TopHat (v1.4.1) and alignment results parsed via SAMtools to generate SAM files. The uniquely and multi-mapped reads were filtered from the BAM files using an MAPQ (mapping quality) of 255 to filter uniquely mapped reads. Read counts to each repeat element were obtained by mapping the multi-mapped reads to each of the repeat classes and counting alignment for each repeat class (all the steps are part of the RepEnrich pipeline). After removing absent repeat elements (zero counts in all samples), the raw counts were then imported to R/Bioconductor package EdgeR differentially expressed genes among samples. The generalized linear model method within the EdgeR package was used to identify differentially expressed genes. We considered genes differentially expressed between two groups of samples when the EdgeR analysis resulted in an adjusted P-value of <0.05.

### 2.8 BODIPY Lipid Staining

#### 2.8.1 Cellular Imaging and Analysis

The day before staining, 10,000 cells were seeded per well in black, clear-bottom, glass 96-well plates pre-coated with 0.01 mg/mL Poly-D-Lysine. The following day, cells were washed once with PBS and fixed with room-temperature 4% paraformaldehyde for 15 minutes. After fixation, cells were washed three times with PBS and incubated with 2 µM Fluorescein isothiocyanate (FITC)-BODIPY 493/503 in PBS for 30 minutes at room temperature in the dark on a rocker. Excess dye was removed by washing the cells three times with PBS.

Cells were then counterstained with 4′,6-diamidino-2-phenylindole (DAPI) (1:10,000) for 5 minutes in the dark, followed by three additional PBS washes. Imaging was performed using a Keyence BZ-X series fluorescence microscope. For each well, six fields were randomly imaged, with a minimum of five wells analyzed per condition per experiment.

Images were analyzed using QuPath Software^40^. Lipid droplets and nuclei were identified using the subcellular detections tool. Detection parameters were adjusted to exclude objects smaller than 0.2 μm^2^ and minimize background artifacts. Total lipid droplet counts were normalized to the number of DAPI-stained nuclei to calculate average droplet count per cell. At least five images per well and five wells per condition were analyzed, using only images with comparable confluency to reduce variability. Image analysis was performed blinded to experimental condition.

#### 2.8.2 Flow Cytometry and Analysis

Following treatment with 48 hour 2’-deoxyadenosine (dAdo; Sigma Aldrich) or 72 hour siDGk, cells were detached with 0.25% trypsin (Thermo Scientific) neutralized, and washed with PBS. Cells were incubated with 2 µM BODIPY 493/503 at 37 °C for 15 minutes protected from light, then washed and resuspended in 200 µL Fluorescence-activated cell sorting (FACS) buffer (PBS, 2 mM EDTA, 2% FBS, 0.05% sodium azide) containing propidium iodide (PI; 1:200; BioLegend). Samples were kept protected from light, on ice, and acquired within 30 minutes on a BD LSR-II Flow Cytometer. BODIPY fluorescence was collected in the FITC channel; PI was collected in a red PI-compatible channel. Data were analyzed in FlowJo v10^41^. Dead cells (PI⁺) were excluded, and BODIPY signal was quantified in the live (PI⁻) gate. Unstained and single-stain compensation controls were included (BODIPY-only cells; PI-only cells generated by heat-killing at 56 °C for 10 minutes), and the compensation matrix was calculated in FlowJo and applied to all samples.

### 2.9 Oil Red O Staining and Quantification

Oil Red O (ORO) staining was performed using the Sigma-Aldrich Lipid (Oil Red O) Staining Kit (Sigma-Aldrich), following the manufacturer’s instructions with modifications described below, to optimize staining for HEPG2 cells.

All steps were performed at 22 °C unless otherwise specified. Cells were fixed in 10% paraformaldehyde (PFA) and incubated for 5 minutes. The PFA was then replaced with fresh 10% PFA, and the cells were incubated for 15 minutes at room temperature on a rocker. Following fixation, the cells were washed with 60% isopropanol and incubated for 1 minute. The cells were allowed to air dry for 10 minutes before the addition of the ORO Working Solution. The cells were then incubated with the ORO Working Solution for 10 minutes at room temperature on a rocker. After incubation, the cells were washed five times with distilled water, covered with water, and imaged using a light microscope.

For quantification, water was removed from the wells, and cells were air-dried in a sterile hood for 10 minutes. 100% isopropanol was then added to each well to elute the ORO stain, followed by a 10 minute incubation at room temperature on a rocker. The solution was mixed thoroughly, transferred to a fresh 96-well plate, and absorbance was measured at 540 nm with background correction at 620 nm using a Spectramax plate reader (Molecular Sciences).

### 2.10 IFN**β** Neutralization Assay

Recombinant human IFNβ neutralizing antibody (nAb; R&D Systems) was added immediately following siRNA transfection at a final concentration of 40 U/mL. Cells were incubated for 72 h post-transfection before RNA isolation, RT-qPCR, and BODIPY lipid staining analyses. Effective IFNβ neutralization was verified by reduced ISG15 mRNA expression as measured by RT-qPCR.

### 2.11 2’-Deoxyadenosine Treatment

2’-deoxyadenosine (dAdo; Sigma Aldrich) was reconstituted to 80 mM in sterile DNase-/RNase-free water per the manufacturer’s instructions. HepG2 cells were seeded into 6- and 24-well plates (2.0 and 0.5 mL media per well, respectively). At 60-70% confluency, cells were treated with EMEM and 10% FBS containing the following dAdo concentrations: 0, 250 µM, 500 µM, or 1 mM (prepared from the 80 mM stock diluted in culture medium, vehicle volume-matched). Plates were maintained at 37 °C, 5% CO₂, and saturated humidity . Cells were harvested at 24 hours for methylation analysis or at 48 hours for RT-qPCR and flow cytometry.

### 2.12 Global DNA methylation (5-mC) ELISA

After 48 hours siRNA transfection or 24 hours dAdo treatment, DNA was isolated from cells using the DNeasy Blood and Tissue Kit (Qiagen). Global 5-mC was quantified using the 5-mC DNA ELISA kit (Zymo Research) with 100 ng genomic DNA per well. DNA was denatured at 98 °C for 5 minutes, chilled on ice for 10 minutes, and the full volume was added to the ELISA plate in a coating buffer (final 100 µL). A 0–25% 5-mC standard curve was run in duplicate on each plate. Absorbance was read at 450 nm. Standard curves were fitted with a four-parameter logistic (4PL; untransformed X) model using 1/Y² weighting, and sample values were interpolated from the fitted curve. Standards/samples outside the calibrated range were excluded or re-assayed after dilution. Reported values are the mean of technical duplicates; where indicated, data are shown as fold change vs control siRNA (siCON).

### 2.13 Statistical analysis

Statistical analyses were performed using Excel (Microsoft 2019) and GraphPad Prism version 9.4.1, GraphPad Software, Inc., San Diego, CA, USA. At least two independent experiments were performed, and error bars represent mean ± standard error of the mean. Two-group comparisons used unpaired, two-tailed Student’s t-tests. For comparisons across ≥3 groups, one-way Analysis of variance (ANOVA) with Dunnett’s multiple-comparisons test was used to compare each treatment to the baseline control. Exact *p-*values are reported in the figure legends.

## 3 Results

### 3.1 Mitochondrial Structure and Function Are Intact in Early DGUOK Deficiency

#### 3.1.1 mtDNA Copy Number

To assess the consequences of early DGUOK depletion in hepatocytes, siRNA-mediated knockdown of DGUOK (siDGk) was performed in HepG2 cells. Knockdown efficiency was confirmed by RT-qPCR after 48 hours, demonstrating 97% reduction in *DGUOK* mRNA expression (p=0.0004), and by western blotting after 72 hours, showing >75% loss of DGUOK protein levels relative to control siRNA (siCON) (**Fig. 1A-B**). To determine whether early DGUOK silencing affects mtDNA, copy number was calculated after 96 hours as a ratio of the mitochondrial genes MT-ND1 and MT-CO1 to the nuclear gene HBB, as previously described^27–29^. mtDNA levels in siDGk cells were comparable to siCON cells, suggesting that prolonged DGUOK deficiency may be required for mtDNA depletion (**Fig. 1C**). Indeed, DGUOK knockout (KO) cells, generated using CRISPR-Cas9 and assessed six weeks after transfection, exhibited >75% reduction in mtDNA copy number compared to the wild-type (WT) cells (p = 0.0034; **Fig. 1C**).

**Figure 1.**
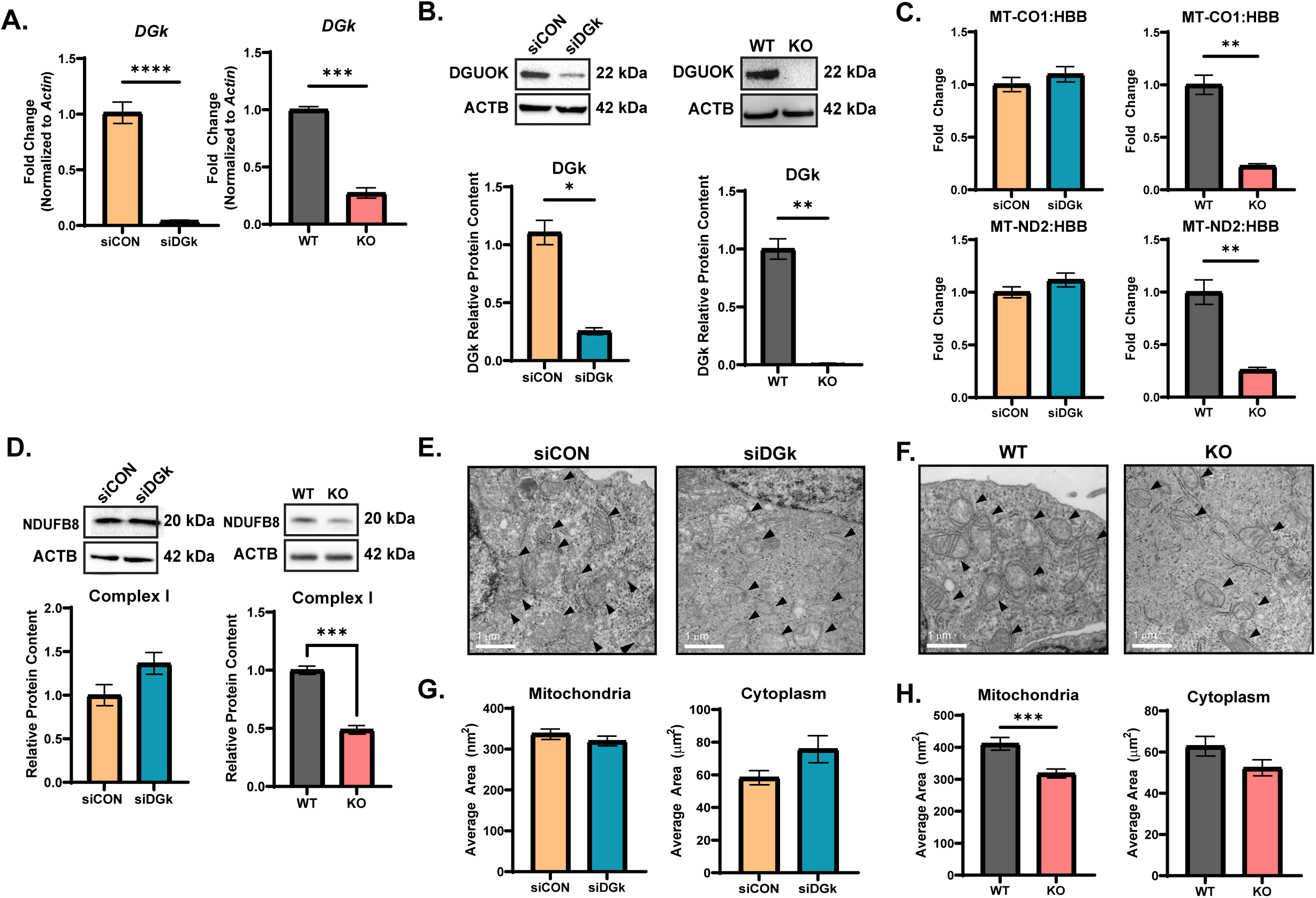
Transient DGUOK Knockdown Maintains mtDNA Integrity and Preserves Mitochondrial Morphology. **(A)** DGUOK mRNA expression was quantified by RT-qPCR in transient siRNA knockdown (siDGk) and CRISPR-Cas9 knockout (KO) HepG2 cells compared to respective controls (siCON, WT). DGUOK expression was reduced by 97% in siDGk (****p = 0.0004, n = 3) and by ∼90% in KO (***p < 0.001, n = 3) **(B)** DGUOK protein levels were assessed by Western blot at 72 h post-transfection or after chronic KO establishment, showing ∼75% loss in siDGk (*p = 0.0457, n = 2) and ∼85% loss in KO (**p = 0.0082, n = 3). **(C)** Mitochondrial DNA (mtDNA) copy number was determined by qPCR using MT-CO1:HBB and MT-ND2:HBB ratios. Transient knockdown showed no significant change (siCON vs siDGk: MT-ND2 p = 0.2986, MT-CO1 p = 0.4273, n = 2–3), while KO cells exhibited significant mtDNA depletion (WT vs KO: MT-ND2 p = 0.0034, MT-CO1 p = 0.0012, n = 3). **(D)** OXPHOS complex analysis showed a 51% reduction in Complex I in KO cells (WT = 1.00, KO = 0.4878, p = 0.006, n = 3), with no changes in Complexes II–V (p > 0.2). siDGk cells maintained normal Complex I levels (p = 0.17, n = 2). (Full OXPHOS panel shown in Fig. S1.) **(E–H)** Transmission electron microscopy (TEM) revealed normal mitochondria (▲) in siDGk cells (E,G), but a 22.6% reduction in mitochondrial cross-sectional area in KO cells (F,H; WT = 411.0 nm², KO = 318.2 nm², p = 0.0001, n = 259–336 mitochondria from 30 cells per group). Cytoplasmic area and mitochondrial occupancy were unchanged between groups (p > 0.05). (Additional quantifications in Fig. S2.). All data represent mean ± SEM; significance was determined by unpaired two-tailed t-tests (*p < 0.05, **p < 0.01, ***p < 0.001, ****p < 0.0001).

#### 3.1.2 Mitochondrial Function and Morphology

Given that mtDNA levels were stable in siDGK cells, mitochondrial integrity was further evaluated to determine whether functional mitochondrial deficits could arise prior to mtDNA depletion. DGUOK deficiency leads to reduced hepatic Oxidative phosphorylation (OXPHOS), with 96% of patients showing specific loss of Complex I protein due to impaired assembly of mtDNA-encoded subunits^9,42^. To assess this phenotype in *DGUOK*-deficient cells, OXPHOS complex proteins were analyzed by Western blot in siDGk and KO cells. DGUOK KO cells exhibited a ∼50% reduction in Complex I protein compared to wild-type cells (p=0.006), while siDGk cells maintained Complex I levels comparable to wild type cells (**Fig. 1D**). Expression of Complexes II–V remained unchanged across both populations (**Fig. S1**).

Next, mitochondrial morphology was examined by transmission electron microscopy (TEM). siDGk cells displayed normal mitochondrial area, whereas DGUOK KO cells showed a 22.6% reduction in mitochondrial cross-sectional area (p=0.0001), consistent with ultrastructural stress and mitochondrial fragmentation reported in mtDNA maintenance disorders^43^ (**Fig. 1E–H**). To confirm that the reduction in mitochondrial area was not the result of reduced cell size, cytoplasmic area and mitochondrial occupancy were quantified and found to be unchanged between groups (**Fig. S2**).

Together, these findings indicate that mitochondrial structure and respiratory function are preserved during early DGUOK depletion, with functional and morphological deficits emerging only upon prolonged loss of DGUOK.

### 3.2 Early DGUOK Deficiency Reprograms Immune, Lipid, and Methylation/Nucleotide Pathways

To investigate gene expression changes following DGUOK knockdown prior to mtDNA depletion, bulk RNA sequencing was performed on siDGk and siCON HepG2 cells. Differential expression analysis identified 2,375 differentially expressed genes (DEGs) between the two groups (padj < 0.10, |log₂FC| ≥ 1.5; **Fig. 2A, S3**). Gene set enrichment analysis (GSEA) using the Reactome database revealed two major categories of transcriptional remodeling: immune signaling pathways were positively enriched in siDGk cells, whereas lipid metabolism pathways were negatively enriched (**Fig. 2B**, **S4–S6**; **Supplementary Tables 1–3**). Immune and antiviral pathways including Interferon Signaling (Normalized enrichment score (NES) = 2.81, False discovery rate (FDR) = 6.9 × 10⁻⁵), Antiviral Mechanisms by IFN-Stimulated Genes (NES = 2.68, FDR = 2.0 × 10⁻⁴), Cytokine Signaling (NES = 5.69, FDR = 0.000), Innate Immune System (NES = 5.57, FDR = 0.000),Viral Infection (NES = 6.86, FDR = 0.000), and JAK/STAT signaling(NES = 2.25, FDR = 0.004) pathways were strongly enriched in siDGk cells (**Fig. 2B, S4–S6; Supplementary Table 4**). Downregulated pathways included Metabolism of Lipids (NES = –8.34, FDR = 0.000), Fatty Acid Metabolism (NES = –5.26, FDR = 0.000), Steroid Metabolism (NES = –5.19, FDR = 0.000), and Cholesterol Biosynthesis (NES = –5.04, FDR = 0.000) (Supplementary Table 4-I). Empirical analysis of digital gene expression data in R (EdgeR) analysis identified 39 human endogenous retrovirus (HERV) elements that were differentially expressed between siDGk and siCON cells (padj < 0.10, |log₂FC| ≥ 1.5; **Fig. 2C–D**). A subset of these HERVs, including members of the ERVK, HERVH, and HERVW families, exhibited moderate to robust levels of upregulation consistent with activation of interferon-associated transcriptional programs^44^.

**Figure 2.**
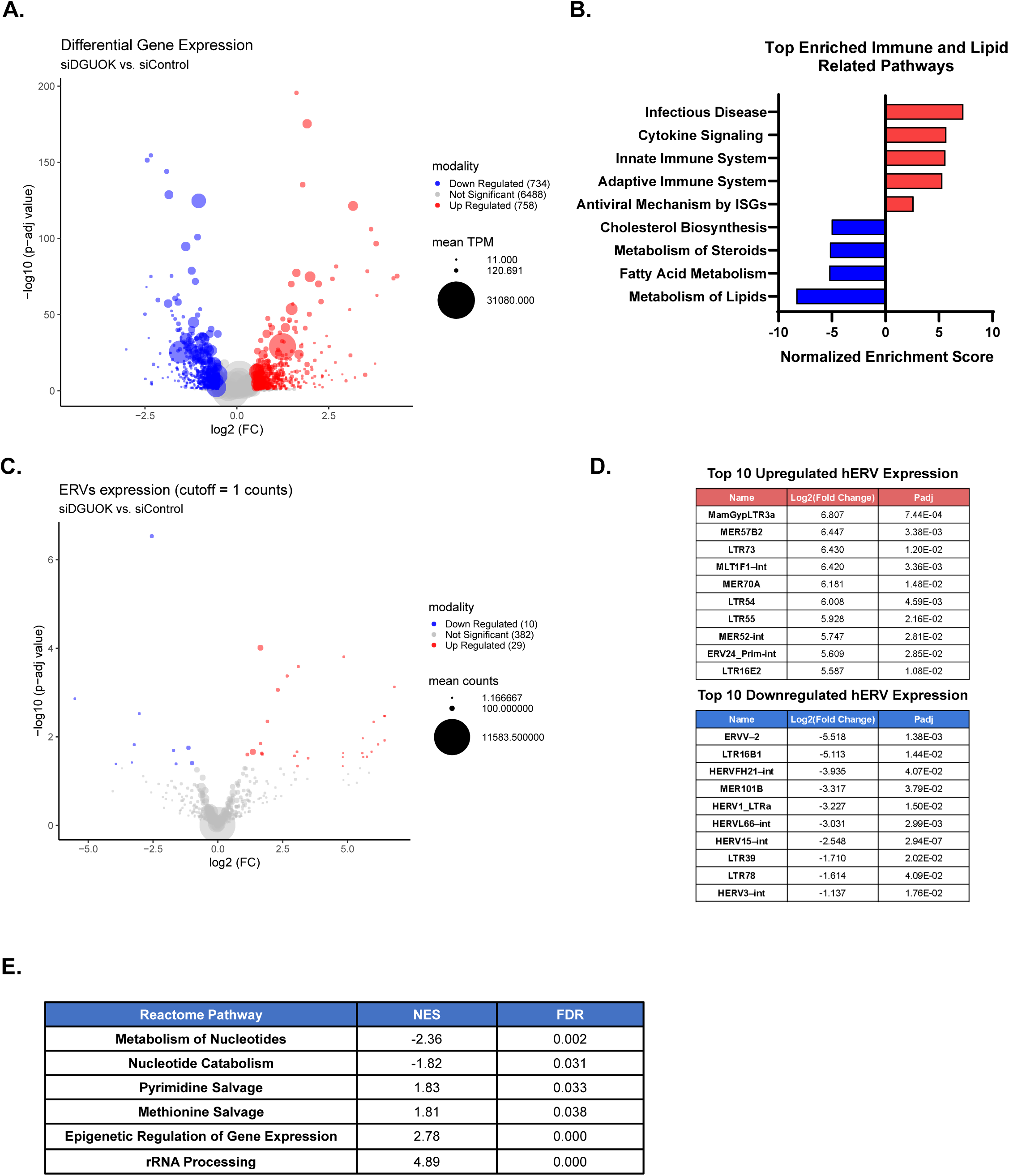
RNA Bulk Sequencing Reveals Positive Enrichment in Innate Immune Pathways and Altered Lipid Metabolism in DGUOK Knockdown Cells. (A) Volcano plot of DEGs between siDGk and siCON HepG2 cells (DESeq2, n = 3 per group). A total of 2,375 genes were significantly altered (padj < 0.10, |log FC| ≥ 1.5), with 758 upregulated and 734 downregulated. (B) Gene set enrichment analysis (GSEA, Reactome) revealed strong upregulation of immune and antiviral pathways, including Interferon Signaling (NES = 2.81, FDR = 6.9 × 10), Antiviral Mechanisms (NES = 2.68, FDR = 2.0 × 10), and Cytokine Signaling (NES = 5.69, FDR = 0.000). Lipid metabolism pathways were conversely suppressed, including Metabolism of Lipids (NES = –8.34) and Fatty Acid Metabolism (NES = –5.26). (C–D) EdgeR analysis of human endogenous retrovirus (hERV) elements identified 39 significantly altered loci (padj < 0.10), including upregulated members of the ERVK, HERVH, and HERVW families linked to interferon-responsive programs (PMID: 36861980). (E) Additional GSEA enrichment included nucleotide and methylation-associated pathways: Metabolism of Nucleotides, Methionine Salvage, and Epigenetic Regulation of Gene Expression.

Expression of the 13 mitochondrially encoded protein-coding genes remained largely unchanged, aside from a modest reduction in MT-CO1 in siDGk cells (**Supplementary Table 2**, **Fig. S7**). This transcript-level stability aligns with preserved mitochondrial protein expression, indicating that transcriptional changes occur before measurable mitochondrial dysfunction.

Pathway enrichment analysis using Reactome highlighted coordinated changes in nucleotide and methyl-donor metabolism (**Fig. 2E**), supporting a model in which DGUOK loss perturbs both purine and methionine metabolic networks. Significantly enriched pathways included Metabolism of Nucleotides (NES = –2.36, FDR = 0.002), Nucleotide Catabolism (NES = –1.82, FDR = 0.031), Pyrimidine Salvage (NES = 1.83, FDR = 0.033), Methionine Salvage (NES = 1.81, FDR = 0.038), Epigenetic Regulation of Gene Expression (NES = 2.78, FDR = 0.000), and rRNA Processing (NES = 4.89, FDR = 0.000). Within these, genes encoding metabolic enzymes central to nucleotide turnover and salvage metabolism, including 5’-nucleotidase, cytosolic II (NT5C2) (log₂FC = –0.58, padj = 1.5 × 10⁻⁴) and purine nucleoside phosphorylase (PNP) (log₂FC = 1.20, padj = 1.4 × 10⁻¹⁷), were differentially expressed along with the core enzymatic components of the methionine cycle such as methionine adenosyltransferase 1A (MAT1A) (log₂FC = –0.78, padj = 7.3 × 10⁻¹⁵) and adenosylhomocysteinase (AHCY) (log₂FC = 0.32, padj = 1.7 × 10⁻⁵).

Collectively, these findings demonstrate for the first time that loss of hepatic DGUOK elicits early, system-wide transcriptional remodeling of innate immune, lipid, and one-carbon metabolism programs preceding mitochondrial perturbations.

### 3.3 DGUOK Deficiency Induces Increased Lipid Accumulation Independently of Mitochondrial Dysfunction

Given transcriptional suppression of lipid metabolism pathways in siDGk, we sought to test whether these changes were reflected at the cellular level. Microvesicular and macrovesicular steatosis are frequently reported in the liver of patients with DGUOK deficiency and accompany progressive liver failure^10,45,46^. To determine whether lipid accumulation occurs in the setting of DGUOK deficiency independently of mtDNA depletion, intracellular lipid droplets were quantified using 4-Difluoro-4-bora-3a,4a-diaza-s-indacene (BODIPY) staining of neutral lipids, which are the core components of lipid droplets. siDGk cells displayed a significant increase in lipid droplet number, averaging 10.29 per cell compared to 6.8 per cell in siCON cells (p=0.0034; **Fig. 3A-B**). This phenotype was corroborated by fluorescence quantification of Oil Red O, another neutral lipid stain, that indicated a 1.75-fold increase of total lipid content in siDGk cells compared to siCON cells (p=0.0001; **Fig. 3C-D**). BODIPY and Oil Red O staining provide limited ultrastructural detail, so lipid morphology was further evaluated using TEM.

**Figure 3.**
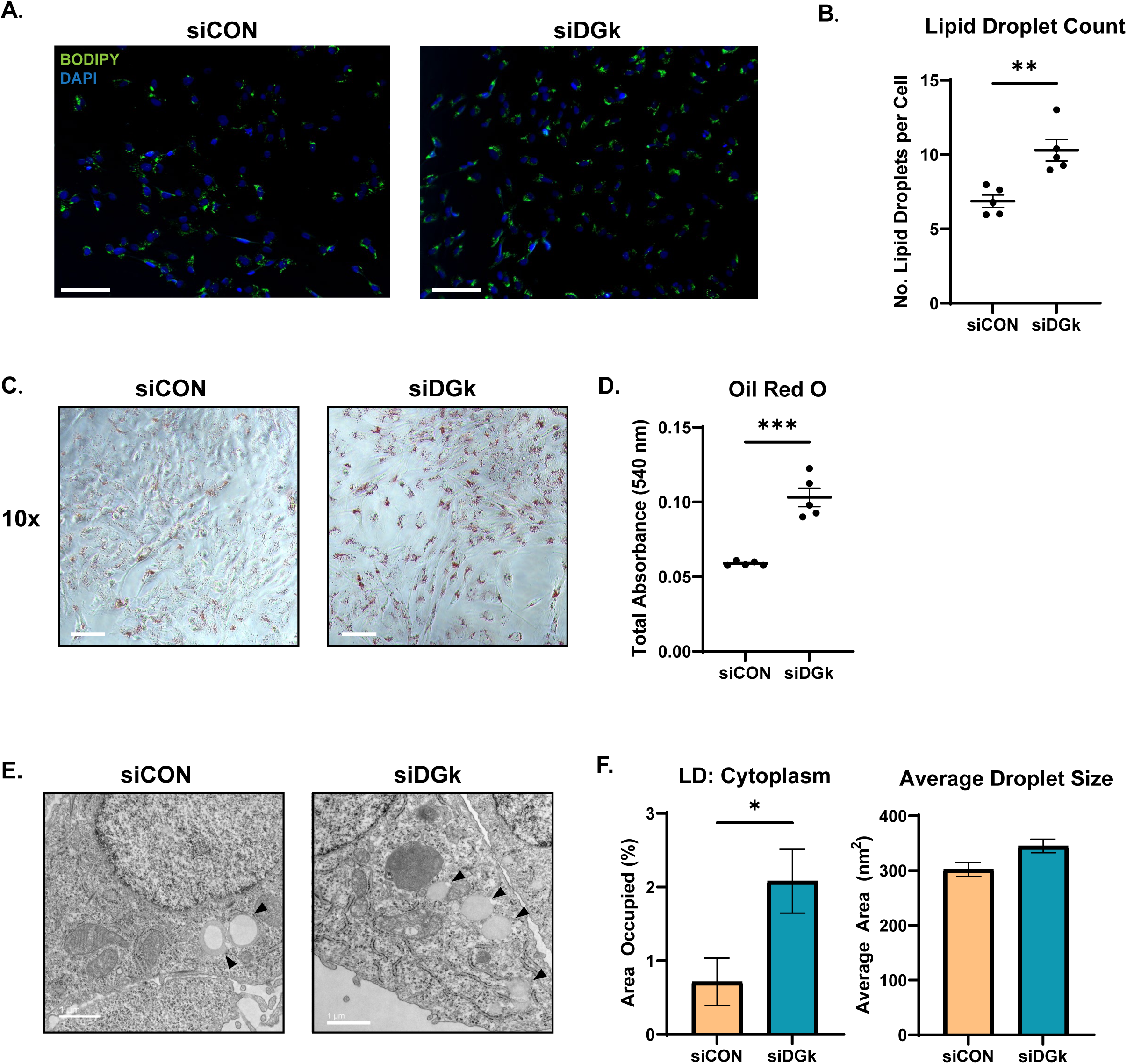
Transient DGUOK knockdown increases lipid accumulation in HEPG2 cells. (A) Neutral lipid staining using BODIPY 493/503 to visualize lipid droplets in HEPG2 cells 72 hours post-transfection with siCON or siDGk. Scale bar = 200 µm. (B) Quantification of BODIPY staining reveals a significant increase in lipid droplet accumulation in siDGk cells compared to siCON (p = 0.0034). Quantification was performed across n = 5 fields of view per condition from 3 independent wells. A minimum of 300 cells were analyzed per condition. (C) Oil Red O staining of neutral lipids in siCON and siDGk cells 72 hours post-transfection. Scale bar = 100 µm. (D) Quantification of Oil Red O staining shows increased lipid accumulation in siDGk cells compared to siCON (p = 0.0001). Quantification was performed across n = 5 wells per condition. (E) Representative transmission electron microscopy (TEM) images showing lipid droplets (▴) in siCON and siDGk HEPG2 cells. (F) TEM quantification reveals significantly increased numbers of cytoplasmic lipid droplets in siDGk cells, with no change in average droplet size (LD:Cytoplasm cell count: n = 30 cells, p = 0.0153; Average droplet size droplet count: siCON n = 313, siDGk n = 344, p = 0.5921). Data are presented as mean ± SEM. Statistical significance was determined using unpaired two-tailed t-tests. ns = not significant.

siDGK cells exhibited a significant 2.9-fold increase in lipid droplet abundance compared to siCON cells (p=0.0153; **Fig. 3E-F**), with mean droplet size slightly elevated (p=0.0167). Together, these findings demonstrate the presence of steatosis in DGUOK-deficient hepatocytes prior to the onset of mitochondrial abnormalities.

### 3.4 Interferon Pathway Blockade Does Not Reverse siDGk–Induced Lipid Accumulation

Type I interferon (IFN-I) signaling has been reported to influence hepatic lipid metabolism, promoting triglyceride accumulation through interferon-stimulated gene (ISG)–dependent remodeling of lipid synthesis and storage^47^. To determine whether the steatotic phenotype observed in DGUOK-deficient hepatocyte cell lines arises secondary to IFN-I activation, we neutralized interferon-β (IFN-β) activity using a monoclonal neutralizing antibody (nAb) in siDGk cells. RT-qPCR verification showed that addition of IFN-β neutralizing antibody significantly reduced ISG15 expression in siDGkcells (p = 0.0051; **Fig. 4A**), indicating that type I interferon signaling was effectively attenuated under these conditions. Despite this reduction, BODIPY staining revealed no improvement in lipid burden following IFN-β blockade, as siDGk+ nAb cells displayed comparable lipid droplet counts to siDGk alone (24.17 ± 2.44 vs. 18.14 ± 2.44 droplets per cell, p = 0.0481; **Fig. 4B–C**). These findings indicate that although DGUOK loss triggers a robust IFN-I response, the accompanying steatosis occurs independently of IFN-β mediated signaling.

**Figure 4.**
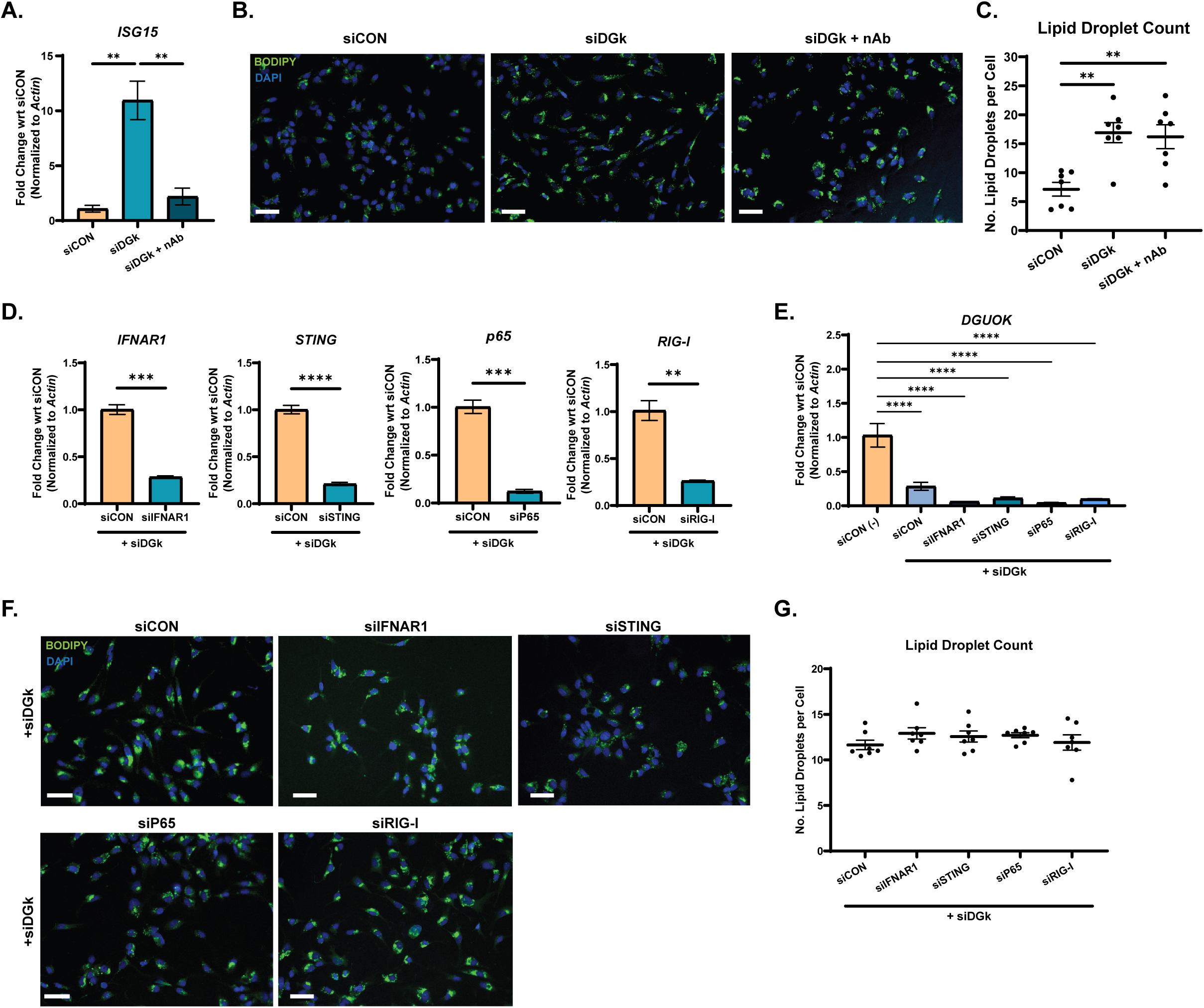
Interferon pathway blockade does not reverse siDGUOK–induced lipid accumulation. **(A)** ISG15 mRNA expression in HepG2 cells transfected with siCON, siDGUOK (siDGk), or siDGUOK + IFNβ neutralizing antibody (nAb) (RT-qPCR, n = 3; normalized to Actb; unpaired two-tailed t-test, p = 0.0051) **(B)** Representative BODIPY (493/503) staining of neutral lipids (green) and DAPI nuclei (blue) 72 h post-transfection. Scale bar = 100 µm. **(C)** Quantification of lipid droplets per cell showing increased lipid accumulation in siDGUOK and siDGUOK + nAb versus siCON (one-way ANOVA with Tukey’s multiple comparisons, n = 7 wells per condition; mean ± SEM). **(D)** Sequential knockdown validation of IFNAR1, STING, p65, and RIG-I by RT-qPCR (cells pre-transfected with innate immune siRNA for 48 h followed by DGUOK silencing for 48 h; n = 3; normalized to Actb; unpaired two-tailed t-tests). **(E)** DGUOK mRNA levels confirming comparable knockdown efficiency across all sequential co-transfection conditions (RT-qPCR, n = 3; one-way ANOVA with Tukey’s test). **(F)** Representative BODIPY/DAPI images of HepG2 cells sequentially transfected with DGUOK and the indicated innate immune siRNAs. Scale bar = 100 µm. **(G)** Quantification of lipid droplets per cell showing that pre-silencing IFNAR1, STING, p65, or RIG-I prior to DGUOK depletion does not alter lipid accumulation (one-way ANOVA with Tukey’s multiple comparisons, n = 7 wells per condition; mean ± SEM). P < 0.05; P < 0.01; P < 0.001; P < 0.0001.

To test whether lipid accumulation reflects activation of upstream innate immune sensors rather than type I IFN signaling itself, cells were sequentially cotransfected with siRNAs targeting DGUOK and one of the core IFN-I pathway mediators. Knockdown of interferon alpha/beta receptor subunit 1 (IFNAR1), stimulator of interferon genes (STING), NF-κB subunit p65 (RELA), and retinoic acid–inducible gene I (RIG-I, DDX58) genes were confirmed by RT-qPCR, with reductions ranging from 72–88% relative to siCON (p < 0.01 for all; **Fig. 4D**).

DGUOK expression remained comparably suppressed across all co-transfection conditions (**Fig. 4E**). Quantification of BODIPY-stained droplets demonstrated that silencing of IFNAR1, STING, p65, or RIG-I failed to diminish lipid accumulation, with mean droplet number remaining unchanged relative to DGUOK knockdown alone (**Fig. 4F–G**). Collectively, these findings indicate that the lipid phenotype persists despite attenuation of interferon activity, raising the possibility that upstream metabolic disturbances, rather than immune signaling, contribute to the immune–lipid remodeling observed in DGUOK-deficient hepatocytes.

### 3.5 Purine Supplementation Recapitulates Lipid and Immune Phenotypes of DGUOK Deficiency

Metabolic disorders involving alterations in purine metabolism, including deficiency of ADA1, ADA2, PNP, SAMHD1 and others, are linked to spontaneous activation of innate immune pathways^48–50^. Similarly, disruption of SAM, AHCY and GNMT enzymes in the methionine pathway alters hepatic lipid metabolism^51,52^. To explore whether an imbalance of purines is sufficient to evoke the immune and lipid-associated cellular responses observed in DGUOK-deficient hepatocytes, HepG2 cells were treated with dAdo, the bio-active purine nucleoside substrate of DGUOK.

To assess global DNA methylation in siDGK and purine-treated cells, 5-mC levels were quantified by ELISA. Both siDGK and dAdo-treated cells showed ∼40% reduction in 5-mC content (p=0.0264 and p=0.0051, respectively; **Fig. 5A**), consistent with diminished methyl-donor availability. Correspondingly, *MAT1A* expression was significantly decreased in both populations (–0.53-fold in dAdo-treated cells, p = 0.0057; –0.78 log₂FC in siDGK; **Fig. 5B–C**). Interferon-stimulated genes (*ISG15*, *ISG20*, and *OAS1*) were upregulated 1.3-2.0-fold (p < 0.05; **Fig. 5B-C**). Flow cytometry quantification of BODIPY staining revealed a dose-dependent increase in lipid accumulation following dAdo treatment, with BODIPY MFI increasing 1.2-fold at 500 μM and 1.4-fold at 1000 μM compared to vehicle controls (p=0.0174 and p=0.0004, respectively; **Fig. 5D**). Similarly, siDGK cells exhibited a 1.3-fold increase in BODIPY MFI relative to siCON (**Fig. 5D**; p=0.0260). Together, these studies show that early DGUOK deficiency and dAdo supplementation elicit parallel alterations in hepatic lipid metabolism, DNA methylation, and interferon signaling in hepatocytes, revealing shared transcriptional and metabolic signatures that emerge independently of mtDNA depletion and mitochondrial dysfunction.

**Figure 5.**
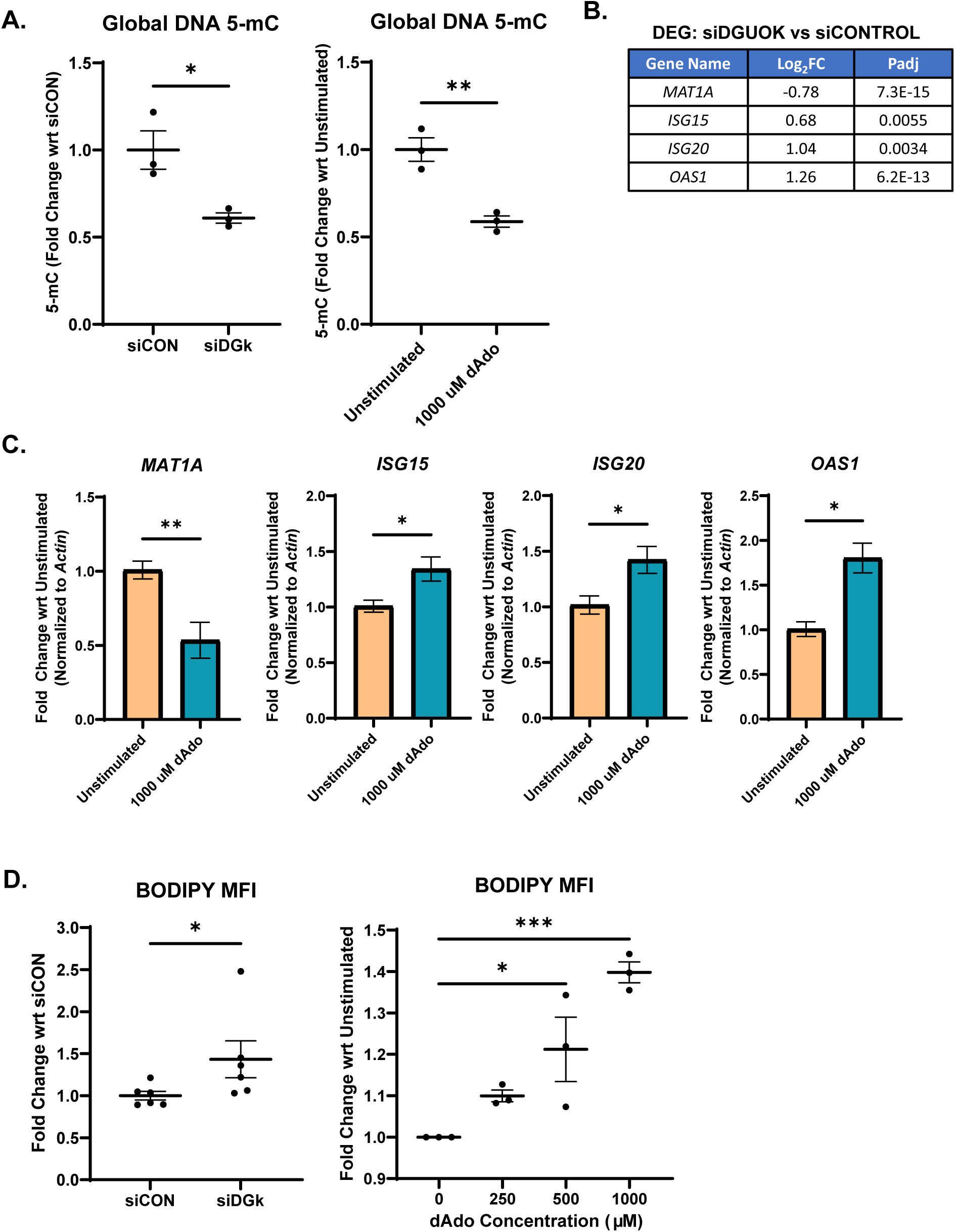
DGUOK depletion and deoxyadenosine exposure reduce DNA methylation, elevate interferon-stimulated gene expression, and increase lipid accumulation. (A) Global DNA 5-mC quantification by ELISA showing reduced methylation in siDGK vs siCON (unpaired two-tailed t-test, p = 0.0264, n = 3) and in dAdo-treated cells (1000 µM) vs unstimulated controls (unpaired two-tailed t-test, p = 0.0051, n = 3). (B) Differentially expressed genes from bulk RNA-seq comparing siDGOK and siCONTROL HepG2 cells, highlighting up-regulated interferon-stimulated genes (ISG15, ISG20, OAS1) and down-regulation of MAT1A involved in methylation metabolism (DESeq2, adjusted p < 0.05). (C) RT-qPCR validation of MAT1A, ISG15, ISG20, and OAS1 expression following dAdo treatment (1000 µM vs unstimulated). Statistical analysis by unpaired two-tailed t-test: MAT1A (p = 0.0057 **), ISG15 (p = 0.0205 *), ISG20 (p = 0.0199 *), and OAS1 (p < 0.05; n = 6 per group). Error bars represent mean ± SEM. (D) Flow cytometric analysis of BODIPY fluorescence intensity showing increased lipid accumulation in both DGUOK-silenced (siDGK) cells and dAdo-treated cells. siDGK vs siCON, Mann–Whitney U test (U = 4, p = 0.0260, n = 6 per group). dAdo dose response, one-way ANOVA with Dunnett’s post-hoc test (F(3,8) = 16.87, p = 0.0008, R² = 0.8635; 500 µM vs 0 µM p = 0.0174 *, 1000 µM vs 0 µM p = 0.0004 ***; n = 3 per group).

## Discussion

This study is the first to identify DGUOK as a regulator of hepatocellular immunometabolism whose function extends beyond mtDNA maintenance. We show that in the short-term, loss of DGUOK in hepatocytes preserves mtDNA copy number, respiration protein expression and mitochondrial architecture yet induces a cell-intrinsic IFN-I/ISG program, accompanied by transcriptional upregulation of multiple species of hERV elements, in addition to profound changes in cellular lipid metabolism and storage, i.e. steatosis. These changes in DGUOK-deficient cells are mirrored by pharmacologic disruption of purine balance (i.e. exogenous exposure to dAdo, the bioactive nucleoside substrate of DGUOK) in wild-type cells, which likewise drives global DNA hypomethylation, ISG induction, and lipid accumulation. By contrast, prolonged DGUOK loss produces the expected mtDNA depletion, Complex I deficiency and mitochondrial ultrastructural abnormalities. Together, these data support a two-stage model in which purine-linked epigenetic remodeling of immune and lipid genes precedes mitochondrial failure in the setting of DGUOK deficiency.

RNA-seq analysis of DGUOK-deficient cells revealed acute and coordinated remodeling of purine, methionine/SAM and methylation-linked transcriptional pathways alongside robust activation of hERVs and ISG **(Fig. 2B-E) (Supplementary Table 3)**. This is best explained by DGUOK-dependent disruption in purine nucleoside salvage metabolism, which has been shown to limit methionine/SAM cycle metabolism and methyl-donor capacity, thereby reducing DNA trans-methylation and de-repressing heavily methylated hERV loci in the genome^6,53,54^ **(Supplementary Table 4M)**. Retrovirus-derived hERV transcripts contain significant secondary structures which form double-stranded RNA-like configurations that engage cellular innate immune receptors including RIG-I/MDA5 and TLR3, stimulating spontaneous IFN-I/ISG signaling and broad innate immune activation^22,44,53,55,56^, precisely the signature observed in DGUOK-deficient cells **(Fig. 2B-D)**. The finding that dAdo supplementation alone is sufficient to phenocopy acute DGUOK loss strengthens the causal link between purine imbalance and viral-mimicry–driven IFN/ISG responses^18,48^. Thus, DGUOK functions as a metabolic checkpoint connecting mitochondrial nucleoside salvage to epigenetic tone and antiviral surveillance.

Concomitant to induction of hERVs and innate immune gene signatures, suppression of lipid metabolism gene programs alongside increased neutral lipid staining and ultrastructural lipid droplet accumulation in siDGUOK cells indicate that steatosis is another early consequence of DGUOK loss, independent of changes in mtDNA copy number **(Fig. 2B, 3B-F) (Supplementary Table 4 and 3I)**. Mechanistically, disrupting IFN-I/ISG signaling had no effect on lipid accumulation **(Fig. 4G)**. In contrast, purine dysregulation and downstream methionine/methylation perturbation appears to drive both immune and lipid phenotypes in hepatocytes **(Fig. 5A-D) (Supplementary Table 4)**. Our observation that dAdo treatment lowers global 5-mC and increases both ISG expression and lipid content in wild-type cells establishes a direct functional bridge from purine imbalance to methyl-donor insufficiency to inflammation and steatosis, independent of mtDNA depletion or respiratory chain collapse **(Fig.5A-D)**. Indeed, purine-mediated reduction of universal methyl-donor SAM availability, which has been described in diverse cellular systems^18,57–59^, provides a unifying explanation for immune and lipid phenotypes. In addition to epigenetic changes, reduced SAM constrains phosphatidylcholine synthesis (PEMT pathway) and very-low-density lipoprotein (VLDL) export, favoring hepatic triglyceride storage^51,60,61^. These data align with clinical reports of steatosis observed in DGUOK-deficient patients even in cases where mtDNA copy number is above the 30% threshold for depletion, positioning DGUOK within a purine-SAM-immune-lipid axis governing hepatocyte homeostasis.

The innate immune activation documented here in DGUOK-deficient hepatocytes echoes signatures previously reported in human deficiencies of ADA1/ADA2 and PNP, as well as dNTP-pool disorders such as SAMHD1-related interferonopathies where purine pathway disruption appears to perturbs methylation and nucleic acid sensing^18,20,48,62–64^. Our data now extend this concept to a mitochondrial nucleoside-salvage enzyme, showing that immunostimulatory consequences can arise upstream of changes in mitochondrial purine metabolism. Furthermore, the delayed emergence of mtDNA deficiency, Complex I loss and mitochondrial fragmentation in DGk-KO cells suggests that purine-driven immunometabolic stress may predispose to, or accelerate, the later bioenergetic failure that typifies MDS diseases.

Current therapeutic strategies for DGUOK deficiency emphasize mtDNA restoration via deoxynucleoside monophosphate supplementation or DGUOK gene replacement in the liver^65,66^. Our new findings suggest that adjunct approaches targeting immunometabolic sequelae should also be considered. These may include methyl-donor support, for example through SAM or precursor supplementation, to restore epigenetic balance, improve phosphatidylcholine synthesis, and reduce lipid accumulation. Interferon-pathway modulation, for example, JAK–STAT inhibitors, to dampen the hERV-driven IFN-I program and downstream inflammation^67–69^.

Finally, purine pathway normalization, potentially via restoring adenosine-deoxyadenosine catabolism or supporting salvage capacity to correct the proximal metabolic trigger inside hepatocytes and other affected cells^50,70^. These strategies could complement liver-directed gene therapy, particularly for patients who present beyond the neonatal window or with multisystem disease, where complete correction of mitochondrial dysfunction is challenging. Furthermore, our observed IFN-I program suggests that gene therapy for DGUOK deficiency may be limited by activation of innate immune antiviral programs that may interfere with AAV-mediated gene delivery^71–73^.

Our conclusions are drawn from human hepatoma cells, which provide mechanistic clarity but do not capture tissue complexity or systemic metabolism at the organism level. We also infer SAM limitation and purine disequilibrium from transcriptomics and 5-mC measurements rather than direct SAM flux assays, which have been performed in ADA2-deficient cells^6,18,54,55,57,59,74–76^.

While siDGUOK cells retain mtDNA copy number and show no overt OXPHOS defect, we cannot fully exclude subtle mtDNA lesions or mitochondrial nucleic acid release that might also contribute to innate sensing pathways innate immune sensors such as cGAS-STING^77–80^. Future work should focus on directly quantifying purine and one-carbon flux and hERV methylation states, test rescue experiments (i.e SAM repletion) for lipid and IFN phenotypes, dissect the nucleic acid sensing hierarchy (RIG-I/MDA5 vs TLR3 vs cGAS–STING) by genetic or pharmacologic inhibition; extend findings to primary human hepatocytes and in vivo models to evaluate steatosis, cytokine profiles, and mitochondrial outcomes in a physiological context^81–84^.

In conclusion, we propose a model in which DGUOK loss-of-function initiates a purine-dependent, epigenetically mediated program that simultaneously activates a cell-intrinsic antiviral program and reconfigures lipid metabolism before mitochondrial failure. This purine-driven axis implicates a unifying framework for inflammation and steatosis observed in DGUOK deficiency, and recasts mitochondrial nucleotide salvage enzymes as gatekeepers of hepatic immune-metabolic homeostasis. Restoring purine and methyl-donor balance and type I interferon signaling emerges as a rational therapeutic path alongside strategies aimed at restoring mtDNA copy number. By demonstrating that DGUOK loss can trigger innate immune activation and steatosis independently of mtDNA depletion, this work expands the conceptual framework of MDS disease from genome maintenance to metabolic-immune regulation, providing new mechanistic insights and potential therapeutic entry points for mitochondrial liver disorders.

## 2 Author Contributions

SS conceptualized and supervised the study and provided overall direction. MC designed and optimized the experimental approaches and performed all cellular and molecular experiments, including siRNA transfections, flow cytometry, ELISA, RT-qPCR, protein and nucleic acid extraction, and imaging analyses. MC was responsible for data analysis, figure preparation, and drafting of the original manuscript. MR assisted with RT-qPCR, DNA and protein collection, imaging analysis and flow cytometry, and contributed to writing, generating supplementary tables, and editing. JJ conducted bioinformatic analyses of bulk RNA sequencing and human endogenous retrovirus (HERV) datasets and generated corresponding figures. AG assisted with experimental assays and manuscript review. NR performed RT-qPCR validation, assisted with figure preparation, and contributed to editing. MV performed Western blot analyses and contributed to manuscript editing. KK assisted in manuscript editing. PS provided technical input during assay development and assisted with flow cytometry setup and manuscript editing.

Funding acquisition and overall project supervision were provided by SS. All authors reviewed, edited, and approved the final manuscript.

## Acknowledgements

The authors thank Guillaume Castillon and Ying Jones of the UC San Diego Electron Microscopy Core (RRID: SCR_022039) for expert assistance with sample preparation, sectioning, and transmission electron microscopy imaging, and Zbigniew Mikulski of the LJI Microscopy Core for training and assistance with lipid imaging. We also acknowledge Jason Greenbaum, Ashmitaa Logandha Ramamoorthy Premlal, Ashu Sethi, and Kai Fung of the LJI Bioinformatics Core for RNA-sequencing analysis support, and Suzanne Alarcon of the LJI Next Generation Sequencing Core for sequencing services. Artem Romanov is thanked for generating the CAS9-KO HepG2 cell line used in this study. Gwen Chan and Christina Doupsas are thanked for their assistance with literature review and sample preparation.

## 4 Data Availability Statement

The Bulk RNA Sequencing and hERV datasets generated for this study can be found at Gene Expression Omnibus under accession number GSE309813(http://www.ncbi.nlm.nih.gov/geo/).

