## Supplemental Figures for "Deoxyguanosine Kinase Deficiency Couples Purine Metabolism to Innate Immune Activation and Lipid Accumulation in Hepatocytes"

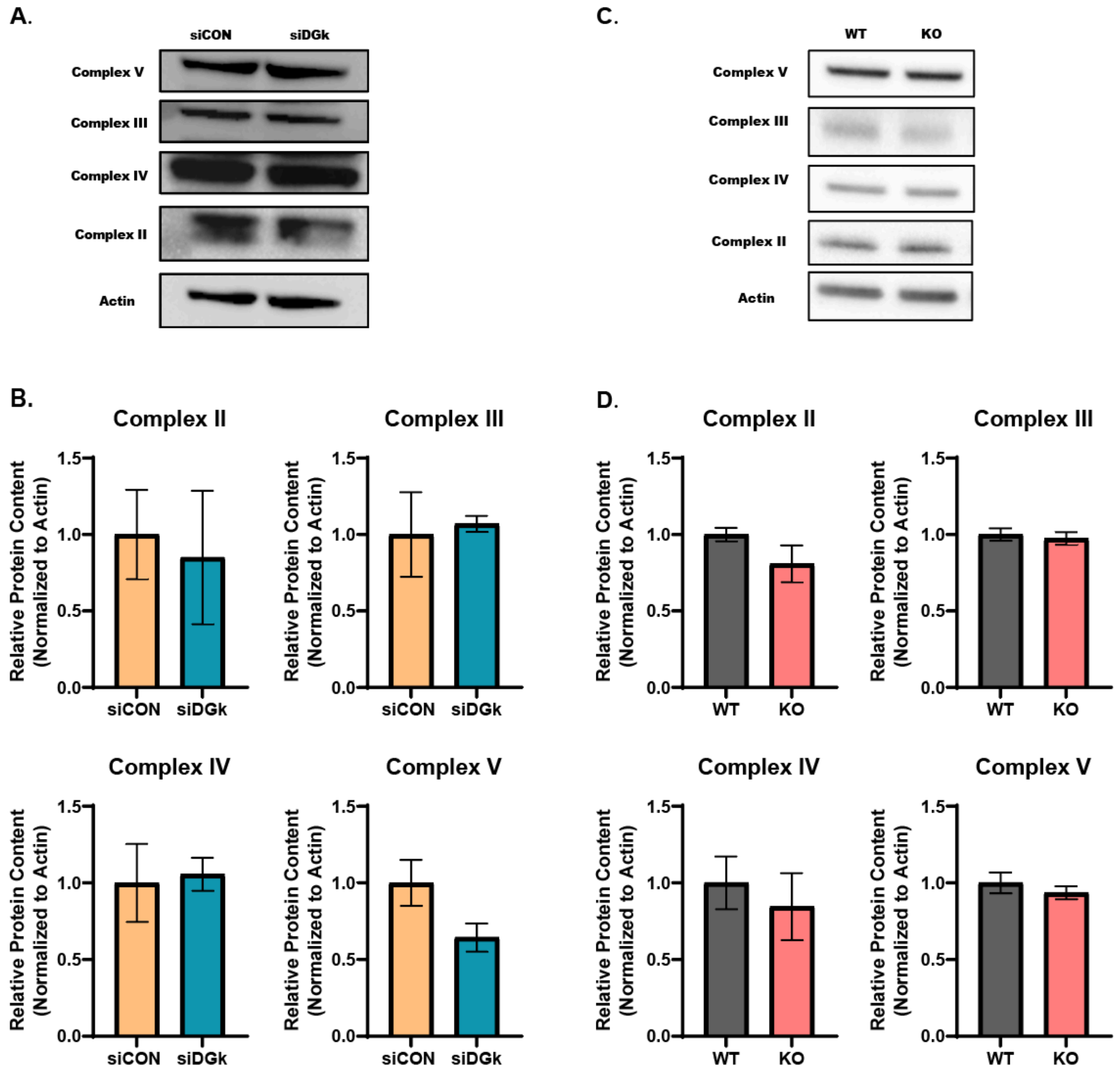


**Figure S1. OXPHOS complex expression in transient and chronic DGUOK-deficient HepG2 cells.** (A–B) Representative Western blots and quantification of mitochondrial oxidative phosphorylation (OXPHOS) complexes II–V in HepG2 cells transfected with siRNA targeting DGUOK (siDGk) or non-targeting control (siCON). Relative protein content was normalized to β-actin. Transient knockdown did not significantly alter the expression of Complex II (siCON = 1.00, siDGk = 0.9953, p = 0.9900), Complex III (siCON = 1.00, siDGk = 0.9442, p = 0.297), Complex IV (siCON = 1.00, siDGk = 0.8629, p = 0.0787), or Complex V (siCON = 1.00, siDGk = 0.8180, p = 0.1932). n = 2 biological replicates per group. (C–D) Representative Western blots and quantification of OXPHOS complexes II–V in DGUOK knockout (KO) HepG2 cells compared to wild-type (WT) controls. No significant changes were observed in Complex II (WT = 1.00, KO = 0.8082, p = 0.209), Complex III (WT = 1.00, KO = 0.9742, p = 0.677), Complex IV (WT = 1.00, KO = 0.8451, p = 0.607), or Complex V (WT = 1.00, KO = 0.9359, p = 0.467). n = 3 biological replicates per group. All data are presented as mean ± SEM and analyzed using unpaired two-tailed t-tests. No statistically significant differences were detected (p > 0.05).

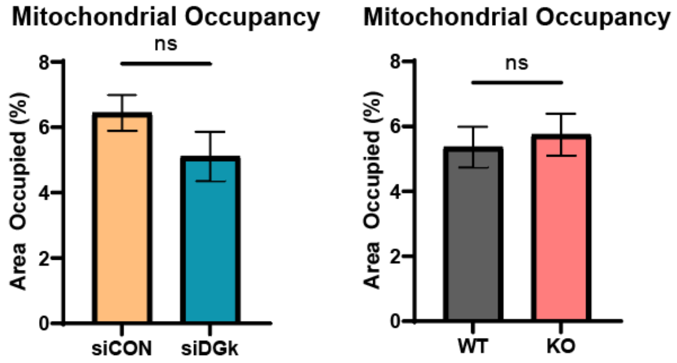


**Figure S2. Transmission electron microscopy (TEM) analysis of mitochondrial occupancy in DGUOK-deficient hepatocytes.** (A) Quantification of mitochondrial occupancy from TEM images of transient (siDGk) and chronic (KO) DGUOK-deficient HepG2 cells compared to controls. Mitochondrial occupancy was calculated as the percentage of cytoplasmic area occupied by mitochondria across 30 randomly selected cells per group. No significant differences were observed between siCON and siDGk cells (siCON = 6.439%, siDGk = 5.108%, $p = 0.1590$, $n = 30$ cells per group) or between WT and KO cells (WT = 5.364%, KO = 5.746%, $p = 0.6727$, $n = 30$ cells per group).

Data are presented as mean ± SEM and analyzed using unpaired two-tailed t-tests. ns, not significant ($p > 0.05$).

A.

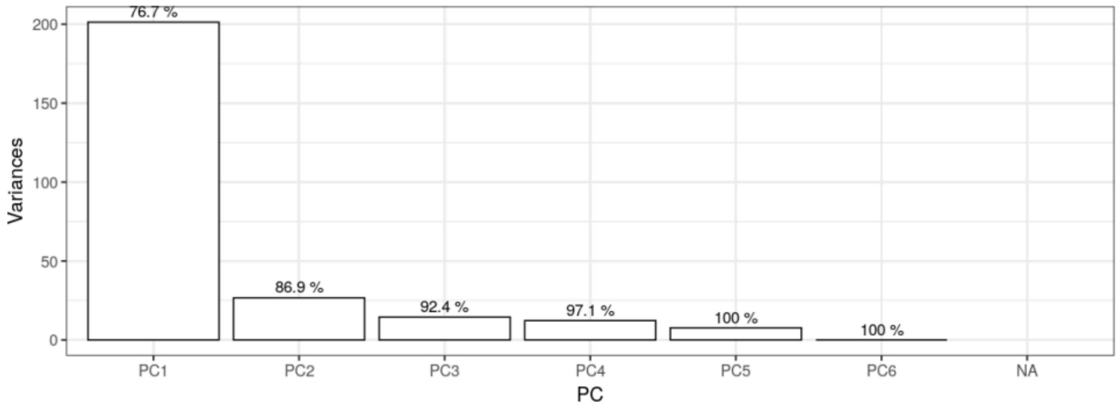


B.

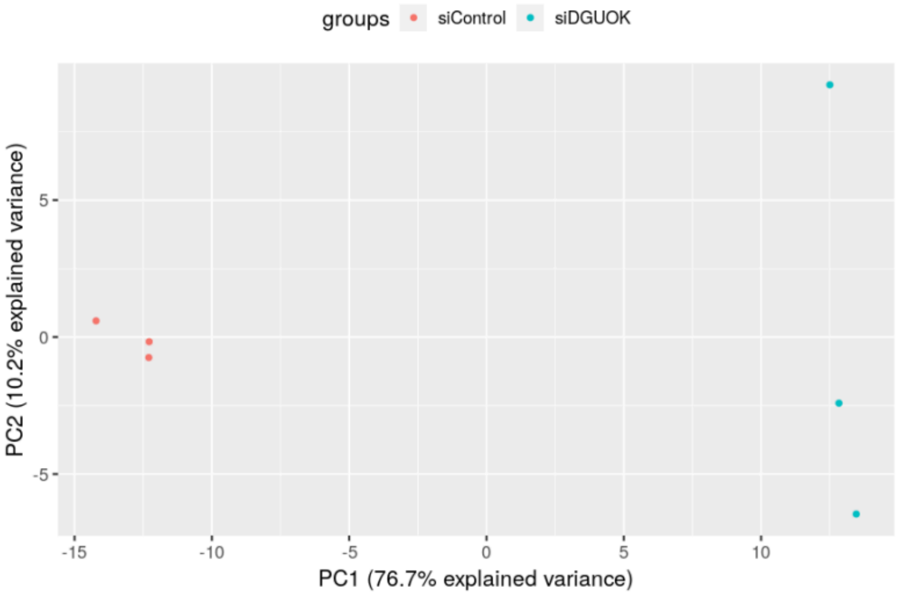


**S3. Principal component analysis (PCA) of RNA-seq data** (A) Scree plot showing the percentage of total variance explained by the rst six principal components. (B) PCA plot of PC1 versus PC2, which together explain 86.9% of the total variance. Samples from siControl and siDGk groups cluster separately, indicating distinct transcriptional proles between conditions.

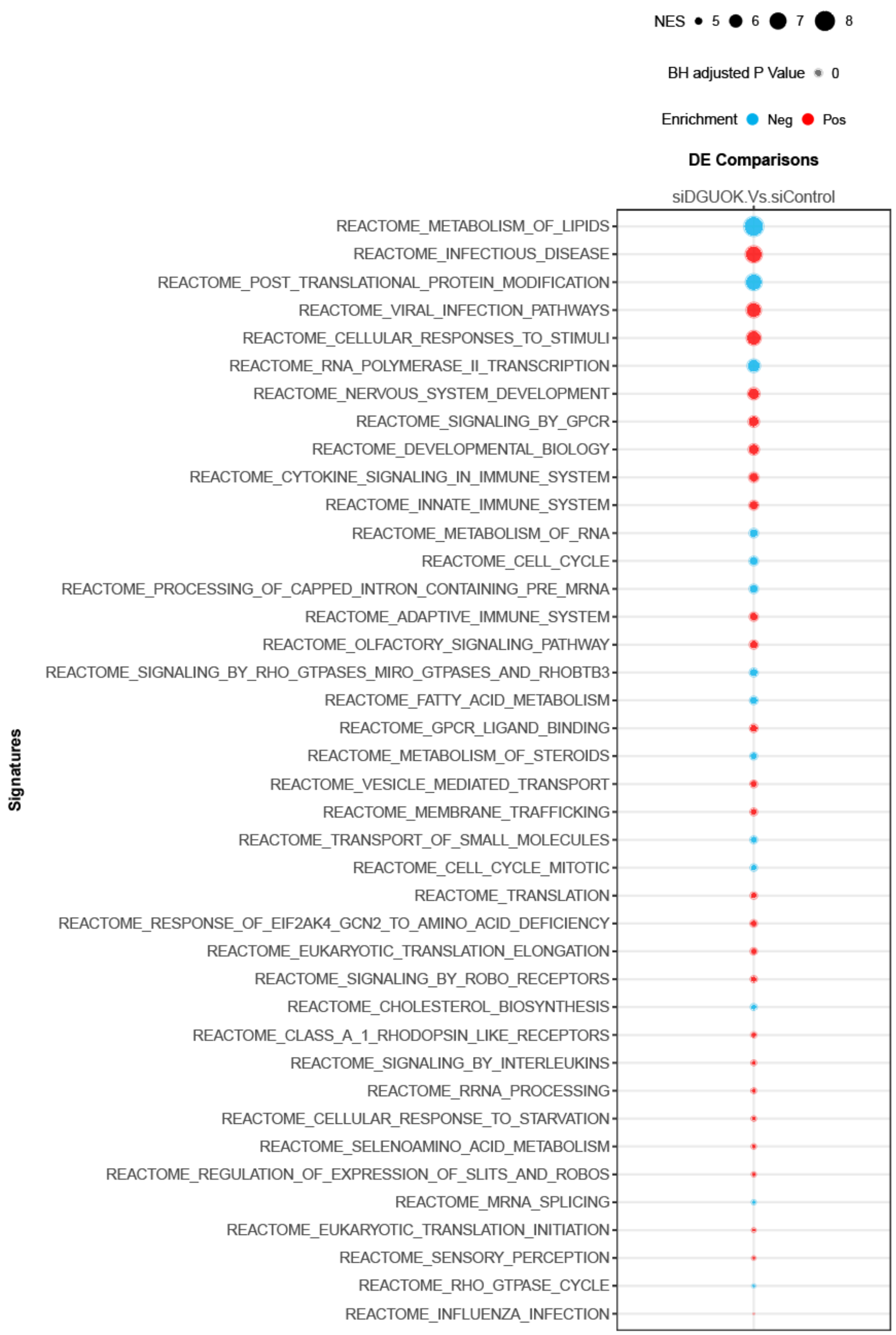


**S4. Reactome Top Enriched Pathways in Gene Enrichment Set Analysis (GSEA).** Dot plot summarizing the most significantly enriched Reactome pathways in siDGk versus siCON HepG2 cells (n = 3 per group). Upregulated immune and antiviral pathways (red) and downregulated lipid-metabolic pathways (blue) are shown with normalized enrichment scores (NES) proportional to dot size. Pathways include Interferon Signaling, Cytokine Signaling, Innate and Adaptive Immune System, Antiviral Mechanisms by IFN-Stimulated Genes, and Metabolism of Lipids, Fatty Acids, Steroids, and Cholesterol.

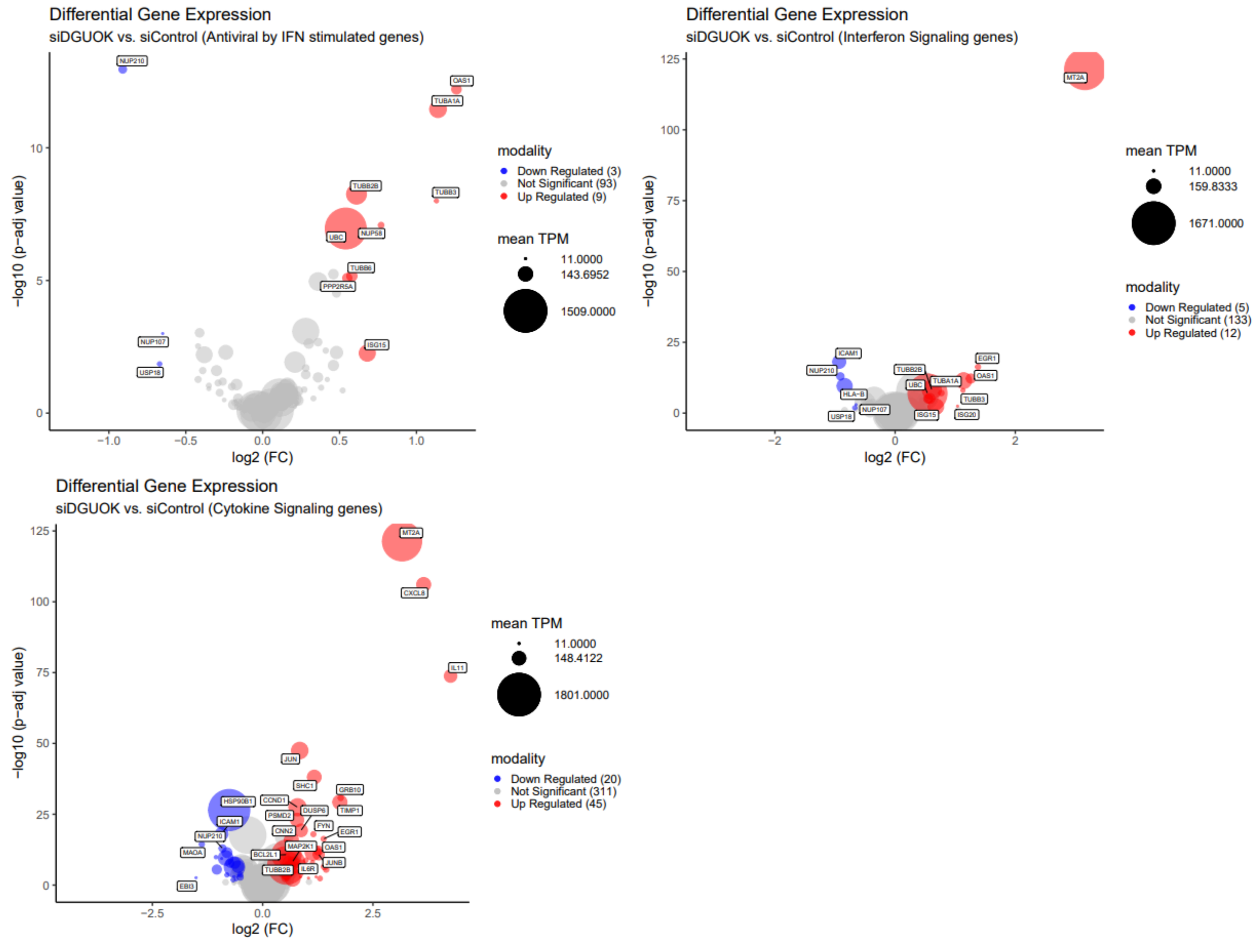


**S5. Differential Gene Expression Volcano Plot of Immune-Related Enriched Pathways.** Volcano plots showing transcriptional changes in antiviral (top), interferon signaling (middle), and cytokine signaling (bottom) gene sets in siDGk versus siCON HepG2 cells (n = 3 per group). Red and blue points represent significantly up- and downregulated genes, respectively (padj < 0.05, $|\log_2 FC| \geq 1.5$). Several interferon-stimulated genes (IFIT1, IFI44L, OAS1, RSAD2) are prominently upregulated, confirming activation of innate immune signaling following DGUOK knockdown.

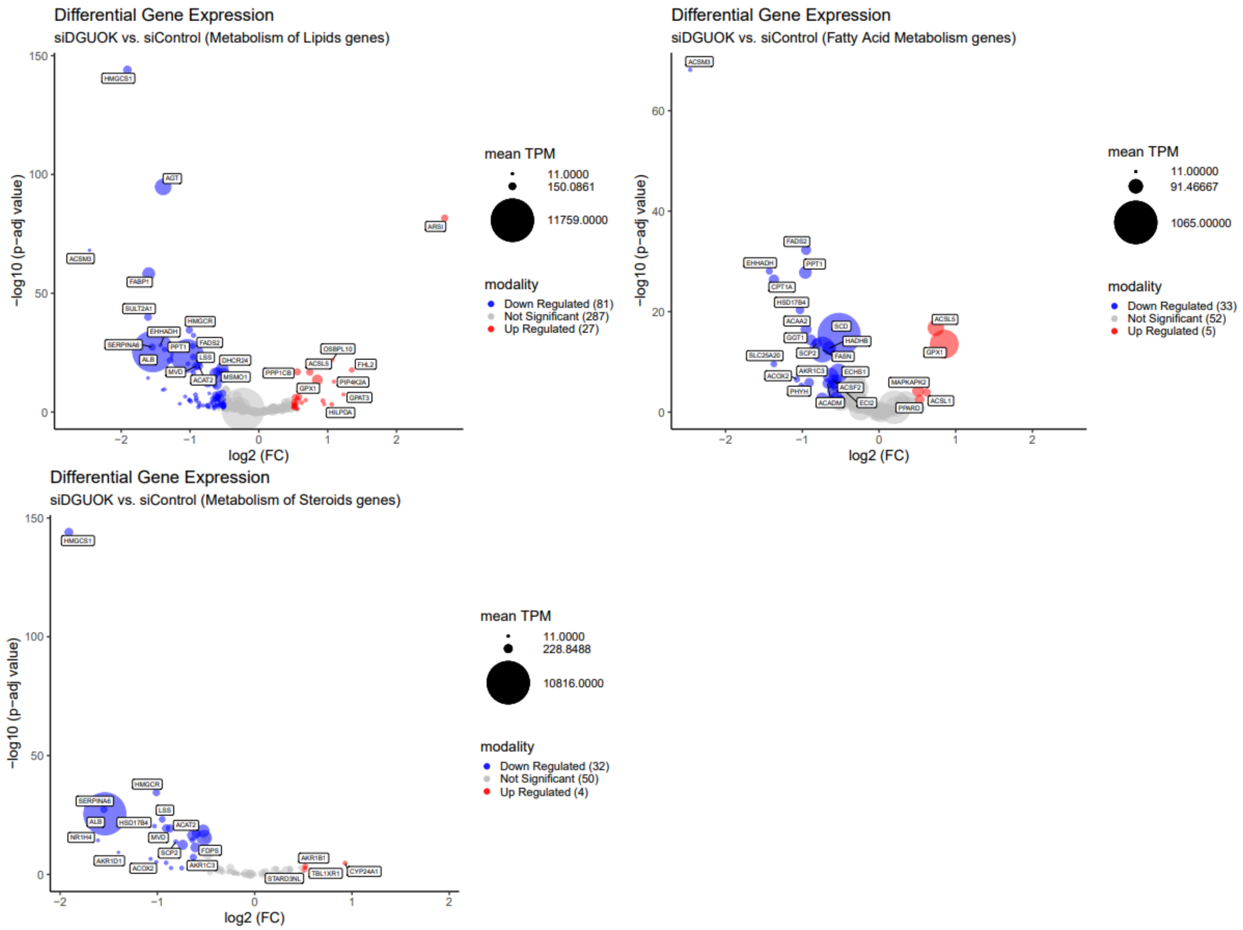


S6. **Lipid Related Plots.** Volcano plots depicting differential expression of genes in (top) Metabolism of Lipids, (middle) Fatty Acid Metabolism, and (bottom) Metabolism of Steroids pathways (n = 3 per group). Blue points indicate significant downregulation of key lipid-metabolic genes including HMGCR, ACACA, FASN, and SCD, consistent with suppressed lipid biosynthesis and β-oxidation programs in siDGk cells.

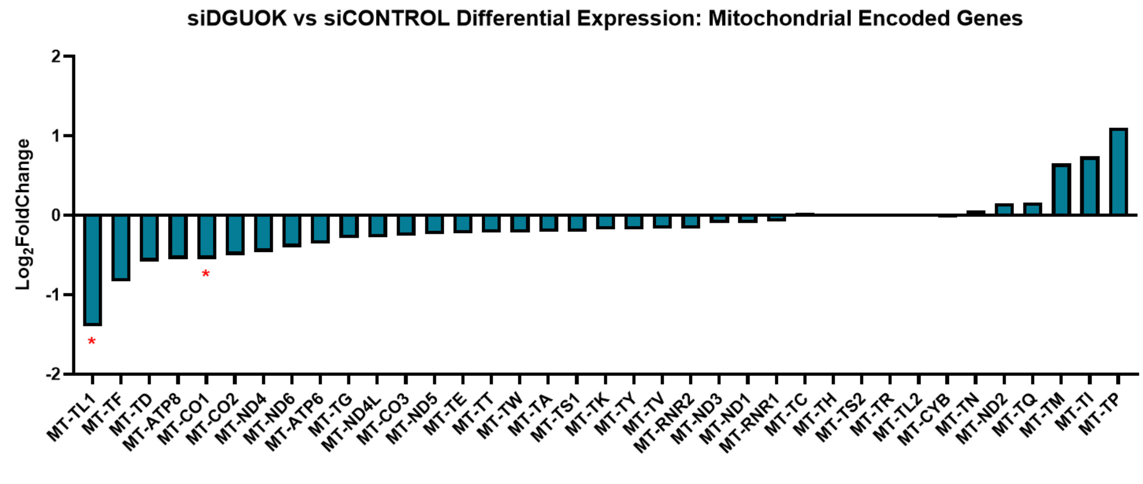


**Figure S7. Mitochondrially encoded gene expression following transient DGUOK knockdown.** Bar plot showing differential expression ($\log_2$ fold change) of mitochondrially encoded genes in siDGk versus siCON HepG2 cells (n = 3 per group). Expression levels were obtained from bulk RNA-seq analysis. Among the 13 protein-coding mitochondrial genes, only protein encoding gene MT-CO1 was significantly downregulated ($\log_2$ FC = –0.56, padj = 0.0044), as well as tRNA gene MT-TL1. All other mitochondrial transcripts, including MT-ND, MT-CYB, and MT-ATP family members, remained unchanged, consistent with preserved mitochondrial mRNA stability at this early stage of DGUOK depletion.
