## Supplemental Tables for "Deoxyguanosine Kinase Deficiency Couples Purine Metabolism to Innate Immune Activation and Lipid Accumulation in Hepatocytes"

| Gene symbol | Full Gene Name | TaqMan probe ID |
|---|---|---|
| *DGUOK* | Deoxyguanosine kinase | Hs00361549_m1 |
| *ACTB* | Actin, beta | Hs01060665_g1 |
| *HBB* | Hemoglobin subunit beta | Hs00758889_s1 |
| *MT-CO1* | Mitochondrially encoded cytochrome c oxidase I | Hs02596864_g1 |
| *MT-ND1* | Mitochondrially encoded NADH dehydrogenase 1 | Hs02596873_s1 |
| *ISG15* | ISG15 ubiquitin-like modifier | Hs01921425_s1 |
| *ISG20* | Interferon-stimulated exonuclease gene 20 | Hs00158122_m1 |
| *MAT1A* | Methionine adenosyltransferase 1A | Hs01547962_m1 |
| *OAS1* | 2'-5'-oligoadenylate synthetase 1 | Hs00973635_m1 |
| *MT-ND2* | Mitochondrially encoded NADH dehydrogenase 2 | Hs02596874_g1 |
| *IFNAR1* | Interferon alpha and beta receptor subunit 1 | Hs01066116_m1 |
| *TMEM173* | Transmembrane protein 173 (STING) | Hs00736955_g1 |

| | | |
|---|---|---|
| *DDX58* | DEAD-box helicase 58 (RIG-I) | Hs01061436_m1 |
| *RELA* | RELA proto-oncogene, NF-κB subunit | Hs01042014_m1 |

**Supplementary Table 1. TaqMan Probes**

This table lists all TaqMan Gene Expression Assays used for quantitative RT-PCR validation of key transcripts identified by RNA-seq in HEPG2 cells.

| **Gene Name** | **Complex** | **$Log_2FC$** | **padj** | **Clinical Depletion in DGUOK Patients** |
|---|---|---|---|---|
| ***MT-ND1*** | I | –0.10 | 0.85 | ↓ mtDNA < 20% |
| ***MT-ND2*** | I | +0.15 | 0.71 | ↓ mtDNA < 20% |
| ***MT-ND3*** | I | –0.10 | 0.53 | ↓ mtDNA < 20% |
| ***MT-ND4*** | I | –0.46 | 0.16 | ↓ mtDNA < 20% |
| ***MT-ND4L*** | I | –0.28 | 0.41 | ↓ mtDNA < 20% |
| ***MT-ND5*** | I | –0.24 | 0.56 | ↓ mtDNA < 20% |
| ***MT-ND6*** | I | –0.40 | 0.36 | ↓ mtDNA < 20% |
| ***MT-CO1*** | IV | –0.56 | 0.0044* | ↓ mtDNA < 20% |
| ***MT-CO2*** | IV | –0.50 | 0.062 | ↓ mtDNA < 20% |
| ***MT-CO3*** | IV | –0.26 | 0.27 | ↓ mtDNA < 20% |
| ***MT-CYB*** | III | +0.01 | 0.99 | ↓ mtDNA < 20% |
| ***MT-ATP6*** | V | –0.36 | 0.35 | ↓ mtDNA < 20% |
| ***MT-ATP8*** | V | –0.55 | 0.13 | ↓ mtDNA < 20% |

**Table 2. Summary of mitochondrially encoded gene expression changes in siDGk versus siCON HepG2 cells and clinical depletion patterns in DGUOK-deficient patients**. Table summarizing $log_2$ fold changes and adjusted p-values for mitochondrially encoded oxidative phosphorylation genes (Complex I–V) from RNA-seq analysis. For clinical comparison, reported mtDNA depletion levels (< 20 %) in DGUOK-deficient patient liver tissue are included(PMID:39079226).

| Gene Symbol | Protein Encoded | Rank in Gene List | Rank Metric Score | Running Enrichment Score | Core Enrichment |
|---|---|---|---|---|---|
| *RCL1* | RNA 3′-phosphate cyclase-like protein (rRNA processing factor) | 147 | 13.9208 | 0.0006 | Yes |
| *RIOK3* | Ser/Thr protein kinase RIO3 | 498 | 5.8539 | -0.0048 | Yes |
| *UTP4* | UTP4/Cirhin, SSU processome component | 563 | 5.2924 | -0.0017 | Yes |
| *RPL10* | 60S ribosomal protein L10 | 619 | 4.8539 | 0.0016 | Yes |
| *RPS11* | 40S ribosomal protein S11 | 684 | 4.4318 | 0.0047 | Yes |
| *RPL22L1* | 60S ribosomal protein L22-like 1 | 685 | 4.4318 | 0.0096 | Yes |
| *RPL31* | 60S ribosomal protein L31 | 883 | 3.6778 | 0.0088 | Yes |
| *RPL7A* | 60S ribosomal protein L7a | 893 | 3.6576 | 0.0135 | Yes |
| *RPS25* | 40S ribosomal protein S25 | 917 | 3.5850 | 0.0178 | Yes |

| | | | | | |
|---|---|---|---|---|---|
| *RPS3A* | 40S ribosomal protein S3A | 940 | 3.5229 | 0.0221 | Yes |
| *XRN2* | 5′→3′ exoribonuclease 2 | 956 | 3.4815 | 0.0266 | Yes |
| *RPL15* | 60S ribosomal protein L15 | 989 | 3.4202 | 0.0306 | Yes |
| *RPL9* | 60S ribosomal protein L9 | 1030 | 3.2518 | 0.0344 | Yes |
| *FCF1* | FCF1, SSU processome factor | 1259 | 2.7212 | 0.0326 | Yes |
| *RPL37* | 60S ribosomal protein L37 | 1392 | 2.4948 | 0.0337 | Yes |
| *RPL26* | 60S ribosomal protein L26 | 1471 | 2.3768 | 0.0363 | Yes |
| *TRMT10C* | tRNA methyltransferase 10C (MRPP1), mitochondrial | 1621 | 2.1805 | 0.0369 | Yes |
| *RPS15A* | 40S ribosomal protein S15a | 1645 | 2.1549 | 0.0412 | Yes |
| *RPL8* | 60S ribosomal protein L8 | 1675 | 2.1249 | 0.0453 | Yes |
| *RPL39* | 60S ribosomal protein L39 | 1706 | 2.0969 | 0.0494 | Yes |

**Supplementary Table 4-A. Reactome "rRNA Processing" — Core Enrichment (Top 20).**

This table lists the top 20 leading-edge genes contributing to enrichment of the Reactome "rRNA Processing" pathway (R-HSA-72312) identified by GSEA preranked analysis comparing siDGUOK and siCON HEPG2 cells. Transient DGUOK knockdown triggered upregulation of nucleolar and ribosome-biogenesis genes including *RCL1*, *UTP4*, and multiple ribosomal protein subunits (*RPL10, RPS11, RPL26*), indicating increased RNA processing and translational turnover prior to mtDNA depletion. Positive rank-metric scores correspond to genes upregulated in the siDGUOK condition, reflecting elevated ribosome biogenesis potentially linked to the observed interferon-stimulated transcriptional state. Functional enrichment within this category aligns with early adaptive remodeling of RNA metabolism secondary to nucleotide imbalance.

| Gene Symbol | Protein Encoded | Rank in Gene List | Rank Metric Score | Running Enrichment Score | Core Enrichment |
| --- | --- | --- | --- | --- | --- |
| *HMGA1* | High mobility group AT-hook 1 (chromatin architectural factor) | 8 | 80.4559 | 0.0011 | Yes |
| *PSMD2* | 26S proteasome regulatory subunit, non-ATPase 2 | 57 | 25.0555 | 0.0009 | Yes |
| *FYN* | Src family tyrosine kinase Fyn | 75 | 19.8539 | 0.0017 | Yes |
| *PPP1CB* | Protein phosphatase 1 catalytic subunit beta (Ser/Thr phosphatase) | 82 | 18.7959 | 0.0029 | Yes |
| *TAF7* | TFIID subunit TAF7 (general transcription factor) | 104 | 17.0915 | 0.0036 | Yes |
| *ITCH* | E3 ubiquitin-protein ligase ITCH | 126 | 14.6778 | 0.0042 | Yes |

| | | | | | |
|---|---|---|---|---|---|
| *TUBA1A* | Tubulin alpha-1A | 157 | 13.1024 | 0.0047 | Yes |
| *AP1S3* | AP-1 adaptor complex subunit sigma-3 | 167 | 12.6576 | 0.0057 | Yes |
| *BCL2L1* | Bcl-xL (BCL2-like 1), anti-apoptotic regulator | 175 | 12.3372 | 0.0068 | Yes |
| *EGFR* | Epidermal growth factor receptor (RTK) | 192 | 11.4949 | 0.0076 | Yes |
| *TUBB* | Tubulin beta chain | 194 | 11.3665 | 0.0089 | Yes |
| *ATP1A1* | $Na^+/K^+$-ATPase alpha-1 subunit | 198 | 11.3098 | 0.0102 | Yes |
| *SH3KBP1* | CIN85 adaptor (endocytosis/signaling) | 224 | 10.4559 | 0.0107 | Yes |
| *IL6R* | Interleukin-6 receptor alpha chain | 229 | 10.2676 | 0.0119 | Yes |
| *SLC25A6* | ADP/ATP translocase 3 (ANT3), mitochondrial carrier | 230 | 10.2596 | 0.0132 | Yes |

| | | | | | |
|---|---|---|---|---|---|
| *TUBB2B* | Tubulin beta-2B | 255 | 9.7212 | 0.0138 | Yes |
| *TUBB3* | Tubulin beta-3 | 269 | 9.4318 | 0.0148 | Yes |
| *SDC1* | Syndecan-1, heparan sulfate proteoglycan | 272 | 9.3279 | 0.0160 | Yes |
| *IRAK1* | IL-1 receptor-associated kinase 1 (Ser/Thr kinase) | 275 | 9.3010 | 0.0173 | Yes |
| *NUP58* | Nucleoporin 58 (Nup58), nuclear pore complex | 311 | 8.4685 | 0.0175 | Yes |

**Supplementary Table 3-B. Reactome "Viral Infection Pathways" — Core Enrichment (Top 20).**

This table summarizes the top 20 leading-edge genes from the Reactome viral infection–related pathways, highlighting strong induction of interferon-responsive and stress-signaling genes in siDGUOK cells. Genes such as *HMGA1*, *EGFR*, *IRAK1*, *IL6R*, and *ITCH* show positive rank-metric scores, consistent with activation of antiviral signaling, cytokine receptor pathways, and proteasome-mediated protein turnover. These transcriptional changes reflect an interferon-stimulated, antiviral-like cellular environment following loss of DGUOK, preceding mitochondrial dysfunction. The enrichment supports that purine disequilibrium activates innate immune programs resembling viral mimicry.

| Gene Symbol | Protein Encoded | Rank in Gene List | Rank Metric Score | Running Enrichment Score | Core Enrichment |
|---|---|---|---|---|---|
| *OSBPL6* | Oxysterol-binding protein-related protein 6 (ORP6) | 28054 | -0.4089 | -0.3501 | Yes |
| *AKR1C2* | Aldo-keto reductase family 1 member C2 (3α-HSD) | 28268 | -0.4318 | -0.3499 | Yes |
| *TGS1* | Trimethylguanosine synthase 1 (cap hypermethylase) | 28430 | -0.4559 | -0.3481 | Yes |
| *ACACB* | Acetyl-CoA carboxylase beta | 28694 | -0.4949 | -0.3493 | Yes |
| *PPARA* | PPAR-α nuclear receptor | 28899 | -0.5229 | -0.3488 | Yes |
| *SEC24B* | COPII coat subunit Sec24B | 29134 | -0.5686 | -0.3491 | Yes |
| *SEC23A* | COPII coat subunit Sec23A | 29206 | -0.5850 | -0.3447 | Yes |
| *PLPP6* | Phospholipid phosphatase 6 | 29259 | -0.5850 | -0.3396 | Yes |
| *FDX1* | Ferredoxin-1 (mitochondrial electron transfer for steroidogenesis) | 29375 | -0.6198 | -0.3365 | Yes |

| | | | | | |
|---|---|---|---|---|---|
| *GC* | Vitamin D-binding protein (group-specific component) | 29683 | -0.6990 | -0.3390 | Yes |
| *LDLRAP1* | LDL receptor adaptor protein 1 | 29854 | -0.7447 | -0.3375 | Yes |
| *CYP7B1* | Oxysterol 7-α-hydroxylase | 29869 | -0.7447 | -0.3313 | Yes |
| *SEC24A* | COPII coat subunit Sec24A | 30560 | -1.0000 | -0.3452 | Yes |
| *AKR1C4* | Aldo-keto reductase family 1 member C4 (3α-HSD) | 30620 | -1.0088 | -0.3404 | Yes |
| *TBL1X* | Transducin beta-like 1, X-linked (coregulator) | 30695 | -1.0458 | -0.3360 | Yes |
| *NFYC* | NF-Y transcription factor subunit gamma | 30716 | -1.0555 | -0.3300 | Yes |
| *CHD9* | Chromodomain helicase DNA-binding protein 9 | 30776 | -1.0809 | -0.3252 | Yes |
| *STARD5* | StAR-related lipid transfer protein 5 | 30786 | -1.0862 | -0.3189 | Yes |
| *CYP17A1* | 17-α-hydroxylase/17,20-lyase | 31108 | -1.2676 | -0.3218 | Yes |

| | | | | | |
|---|---|---|---|---|---|
| *LBR* | Lamin B receptor (sterol-reductase–like; inner nuclear membrane) | 31178 | -1.3188 | -0.3173 | Yes |

**Supplementary Table 4-C. Reactome "Metabolism of Steroids" — Core Enrichment (Top 20).**

The "Metabolism of Steroids" pathway exhibited strong negative enrichment, with leading-edge genes including *AKR1C2*, *FDX1*, *CYP7B1*, and *CYP17A1* downregulated in siDGUOK cells. These genes govern cholesterol, bile acid, and steroid hormone synthesis, processes typically dependent on adequate mitochondrial function and acetyl-CoA availability. Downregulation of these genes parallels the lipidomic and morphological evidence of impaired fatty acid oxidation and reduced steroidogenic activity described in the results. Together, these findings suggest that loss of DGUOK disrupts hepatocellular sterol homeostasis independently of mtDNA depletion.

| Gene Symbol | Protein Encoded | Rank in Gene List | Rank Metric Score | Running Enrichment Score | Core Enrichment |
|---|---|---|---|---|---|
| *ARNT* | Aryl hydrocarbon receptor nuclear translocator (HIF-1β) | 28262 | -0.4318 | -0.2743 | Yes |
| *AKR1C2* | Aldo-keto reductase 1C2 (3α-HSD) | 28268 | -0.4318 | -0.2731 | Yes |
| *LTC4S* | Leukotriene C4 synthase | 28289 | -0.4318 | -0.2723 | Yes |
| *PIK3C2G* | Class II PI3K, catalytic subunit C2-γ | 28341 | -0.4437 | -0.2725 | Yes |
| *GBA3* | Cytosolic β-glucosidase | 28382 | -0.4437 | -0.2723 | Yes |
| *PRKAG2* | AMPK γ2 regulatory subunit | 28411 | -0.4559 | -0.2718 | Yes |
| *MTMR4* | Myotubularin-related phosphatase 4 | 28413 | -0.4559 | -0.2704 | Yes |
| *TGS1* | Trimethylguanosine synthase 1 | 28430 | -0.4559 | -0.2695 | Yes |

| | | | | | |
|---|---|---|---|---|---|
| *ACSBG1* | Very-long-chain acyl-CoA synthetase (bubblegum 1) | 28520 | -0.4685 | -0.2708 | Yes |
| *PLA2G5* | Group V secreted phospholipase A2 | 28529 | -0.4685 | -0.2697 | Yes |
| *CYP4B1* | Cytochrome P450 4B1 (ω-hydroxylase) | 28541 | -0.4685 | -0.2686 | Yes |
| *FUT1* | Fucosyltransferase 1 (H-antigen synthesis) | 28556 | -0.4685 | -0.2677 | Yes |
| *PIK3C2B* | Class II PI3K, catalytic subunit C2-β | 28623 | -0.4815 | -0.2683 | Yes |
| *PTEN* | PTEN lipid/protein phosphatase (PI3K/Akt brake) | 28646 | -0.4815 | -0.2675 | Yes |
| *RUFY1* | RUN and FYVE domain-containing protein 1 (endosomal trafficking) | 28648 | -0.4815 | -0.2662 | Yes |
| *TNFAIP8L1* | TNFAIP8-like 1 (TiPEL1; | 28654 | -0.4815 | -0.2649 | Yes |

| | | | | | |
|---|---|---|---|---|---|
| | lipid-binding/apoptosis) | | | | |
| *ACACB* | Acetyl-CoA carboxylase β | 28694 | -0.4949 | -0.2647 | Yes |
| *SMPD4* | Neutral sphingomyelinase 3/4 (sphingomyelin phosphodiesterase) | 28709 | -0.4949 | -0.2638 | Yes |
| *ACSL6* | Long-chain acyl-CoA synthetase 6 | 28719 | -0.4949 | -0.2627 | Yes |
| *ACOT6* | Acyl-CoA thioesterase 6 | 28736 | -0.4949 | -0.2618 | Yes |

**Supplementary Table 4-D. Reactome "Metabolism of Lipids" — Core Enrichment (Top 20).**

This table details the top 20 downregulated genes driving negative enrichment of the "Metabolism of Lipids" pathway. Core genes such as *PPARA*, *ACACB*, *PLA2G5*, *PTEN*, and *PRKAG2* exhibited negative rank-metric scores, consistent with suppression of fatty acid β-oxidation, phospholipid remodeling, and peroxisomal transport. The collective downregulation of lipid metabolic genes supports the accumulation of intracellular lipids observed by BODIPY and Oil Red O staining in siDGUOK HEPG2 cells and reinforces that metabolic reprogramming occurs prior to measurable mitochondrial genome loss.

| Gene Symbol | Protein Encoded | Rank in Gene List | Rank Metric Score | Running Enrichment Score | Core Enrichment |
|---|---|---|---|---|---|
| *MT2A* | Metallothionein-2 A (metal/oxidative-stress binder) | 3 | 124.6383 | 0.0039 | Yes |
| *EGR1* | Early growth response 1 transcription factor | 91 | 18.1427 | 0.0054 | Yes |
| *OAS1* | 2′-5′-oligoadenylate synthetase 1 (RNase L pathway) | 148 | 13.8861 | 0.0077 | Yes |
| *TUBA1A* | α-Tubulin 1A (microtubules) | 157 | 13.1024 | 0.0115 | Yes |
| *TUBB2B* | β-Tubulin 2B | 255 | 9.7212 | 0.0126 | Yes |
| *TUBB3* | β-Tubulin 3 | 269 | 9.4318 | 0.0162 | Yes |
| *YBX1* | Y-box binding protein 1 (mRNA/DNA binding) | 285 | 8.9586 | 0.0198 | Yes |
| *NUP58* | Nucleoporin 58 (nuclear pore) | 311 | 8.4685 | 0.0231 | Yes |

| | | | | | |
|---|---|---|---|---|---|
| *UBC* | Polyubiquitin C (ubiquitin precursor) | 322 | 8.3468 | 0.0268 | Yes |
| *PLCG1* | Phospholipase C-γ1 (signal transduction) | 435 | 6.4685 | 0.0275 | Yes |
| *TUBB6* | β-Tubulin 6 | 444 | 6.3979 | 0.0313 | Yes |
| *PPP2R5A* | PP2A regulatory subunit B56-α | 452 | 6.3188 | 0.0351 | Yes |
| *TUBA1C* | α-Tubulin 1C | 469 | 6.1739 | 0.0386 | Yes |
| *NUP43* | Nucleoporin 43 | 519 | 5.6990 | 0.0412 | Yes |
| *HSPA2* | HSP70 family chaperone (HSPA2) | 707 | 4.3098 | 0.0396 | Yes |
| *TUBA1B* | α-Tubulin 1B | 772 | 4.0915 | 0.0417 | Yes |
| *UBE2I* | UBC9 — SUMO-conjugating enzyme E2 | 905 | 3.6198 | 0.0418 | Yes |
| *PPP2CA* | PP2A catalytic subunit α | 930 | 3.5528 | 0.0451 | Yes |

| | | | | | |
|---|---|---|---|---|---|
| *TRIM8* | TRIM8 E3 ubiquitin ligase | 948 | 3.5086 | 0.0486 | Yes |
| *ISG20* | Interferon-stimulated 3′–5′ exonuclease (antiviral) | 992 | 3.3979 | 0.0514 | Yes |

**Supplementary Table 3-E. Reactome “Interferon Signaling” — Core Enrichment (Top 20).** Genes listed in this table represent the top 20 contributors to the positive enrichment of the “Interferon Signaling” pathway, one of the most significantly upregulated categories in DGUOK-deficient cells (NES = 2.81, FDR = $6.9\times10^{-5}$). Upregulated antiviral effectors (*ISG15, ISG20, OAS1*), signaling mediators (*PLCG1, TRIM8*), and cytoskeletal regulators (*TUBA1A, TUBB2B, TUBB3*) indicate coordinated activation of type I interferon response, cytoskeletal remodeling, and nuclear–cytoplasmic trafficking. These findings correspond with increased ISG mRNA expression validated by RT-qPCR and support a link between disrupted purine metabolism and innate immune activation.

| Gene Symbol | Protein Encoded | Rank in Gene List | Rank Metric Score | Running Enrichment Score | Core Enrichment |
|---|---|---|---|---|---|
| *HMGA1* | High mobility group AT-hook 1 (chromatin architectural protein) | 8 | 80.4559 | 0.0008 | Yes |
| *JUN* | c-Jun, AP-1 transcription factor | 21 | 50.0915 | 0.0015 | Yes |
| *PSMD2* | 26S proteasome non-ATPase subunit 2 | 57 | 25.0555 | 0.0015 | Yes |
| *CTNNB1* | β-Catenin (Wnt effector/adherens junctions) | 67 | 21.5850 | 0.0023 | Yes |
| *FYN* | Src-family tyrosine kinase Fyn | 75 | 19.8539 | 0.0031 | Yes |
| *PPP1CB* | Protein phosphatase 1 catalytic subunit β | 82 | 18.7959 | 0.0040 | Yes |
| *HGS* | HRS/HGS, ESCRT-0 endosomal adaptor | 96 | 17.7212 | 0.0047 | Yes |

| | | | | | |
|---|---|---|---|---|---|
| *TAF7* | TFIID subunit TAF7 | 104 | 17.0915 | 0.0055 | Yes |
| *ACTR3* | ARP3, Arp2/3 actin nucleator | 123 | 15.0506 | 0.0060 | Yes |
| *ACTG1* | γ-Actin (cytoskeleton) | 124 | 15.0269 | 0.0071 | Yes |
| *ITCH* | ITCH E3 ubiquitin-protein ligase | 126 | 14.6778 | 0.0081 | Yes |
| *TUBA1A* | α-Tubulin 1A (microtubules) | 157 | 13.1024 | 0.0083 | Yes |
| *AP1S3* | AP-1 complex subunit σ-3 (vesicle trafficking) | 167 | 12.6576 | 0.0090 | Yes |
| *BCL2L1* | Bcl-xL, anti-apoptotic regulator | 175 | 12.3372 | 0.0099 | Yes |
| *EGFR* | Epidermal growth factor receptor (RTK) | 192 | 11.4949 | 0.0105 | Yes |
| *TUBB* | β-Tubulin (class I) | 194 | 11.3665 | 0.0115 | Yes |

| | | | | | |
|---|---|---|---|---|---|
| *ATP1A1* | $Na^+/K^+$-ATPase α1 catalytic subunit | 198 | 11.3098 | 0.0125 | Yes |
| *CD9* | CD9 tetraspanin (membrane organizer) | 199 | 11.2676 | 0.0135 | Yes |
| *ITPR3* | IP3 receptor type 3 (ER $Ca^{2+}$ channel) | 222 | 10.4815 | 0.0139 | Yes |
| *SH3KBP1* | CIN85/SH3KBP1 adaptor protein | 224 | 10.4559 | 0.0149 | Yes |

**Supplementary Table 3-F. Reactome "Infectious Disease" — Core Enrichment (Top 20).**

This table presents genes from Reactome's "Infectious Disease" module that contribute to positive enrichment in siDGUOK cells. Leading-edge genes such as *JUN*, *CTNNB1*, *ITCH*, *FYN*, and *EGFR* reflect broad activation of host–pathogen defense programs and endosomal trafficking networks. The induction of signaling adaptors (*SH3KBP1, AP1S3*), cytoskeletal components (*ACTR3, ACTG1*), and apoptosis regulators (*BCL2L1*) collectively suggests activation of stress-responsive pathways and cellular remodeling consistent with the interferon-stimulated transcriptional signature.

| Gene Symbol | Protein Encoded | Rank in Gene List | Rank Metric Score | Running Enrichment Score | Core Enrichment |
|---|---|---|---|---|---|
| *LTC4S* | Leukotriene C4 synthase (glutathione-dependent) | 28289 | -0.4318 | -0.3404 | Yes |
| *PRKAG2* | AMPK γ2 regulatory subunit | 28411 | -0.4559 | -0.3381 | Yes |
| *ACSBG1* | Very-long-chain acyl-CoA synthetase (bubblegum 1) | 28520 | -0.4685 | -0.3354 | Yes |
| *CYP4B1* | Cytochrome P450 4B1 (ω-hydroxylase) | 28541 | -0.4685 | -0.3301 | Yes |

| | | | | | |
|---|---|---|---|---|---|
| *ACACB* | Acetyl-CoA carboxylase β (ACC2; FA oxidation control) | 28694 | -0.4949 | -0.3287 | Yes |
| *ACSL6* | Long-chain acyl-CoA synthetase 6 | 28719 | -0.4949 | -0.3235 | Yes |
| *ACOT6* | Acyl-CoA thioesterase 6 | 28736 | -0.4949 | -0.3180 | Yes |
| *PPT2* | Palmitoyl-protein thioesterase 2 (lysosomal) | 28829 | -0.5086 | -0.3148 | Yes |
| *HPGD* | 15-hydroxyprostag landin dehydrogenase | 29048 | -0.5528 | -0.3154 | Yes |
| *SLC27A3* | FATP3, fatty-acid transport protein/acyl-CoA synthetase | 29136 | -0.5686 | -0.3121 | Yes |
| *SLC25A17* | Peroxisomal transporter PMP34 (CoA/FAD carriers) | 29290 | -0.6021 | -0.3107 | Yes |
| *ACOT7* | Acyl-CoA thioesterase 7 | 29494 | -0.6576 | -0.3108 | Yes |

| | | | | | |
|---|---|---|---|---|---|
| *ELOVL7* | Very-long-chain fatty-acid elongase 7 | 29534 | -0.6576 | -0.3060 | Yes |
| *PCTP* | Phosphatidylcholine transfer protein (StARD2) | 29604 | -0.6778 | -0.3022 | Yes |
| *HACD2* | 3-hydroxyacyl-CoA dehydratase 2 (elongation) | 29798 | -0.7212 | -0.3020 | Yes |
| *MCAT* | Malonyl-CoA–ACP transacylase (mitochondrial FAS II) | 30269 | -0.8861 | -0.3100 | Yes |
| *MLYCD* | Malonyl-CoA decarboxylase (regulates FA oxidation) | 30273 | -0.8861 | -0.3042 | Yes |
| *CYP1A1* | Cytochrome P450 1A1 | 30391 | -0.9208 | -0.3017 | Yes |
| *CYP4F11* | Cytochrome P450 4F11 (ω-hydroxylation of eicosanoids) | 30618 | -1.0088 | -0.3025 | Yes |
| *FAAH2* | Fatty-acid amide hydrolase 2 | 30765 | -1.0757 | -0.3010 | Yes |

**Supplementary Table 4-G. Reactome "Fatty Acid Metabolism" — Core Enrichment (Top 20).**

The "Fatty Acid Metabolism" pathway was negatively enriched, with leading-edge genes including *ACACB*, *ACSL6*, *ACOT6*, *ELOVL7*, and *FAAH2* showing decreased expression in siDGUOK cells. These genes govern β-oxidation, elongation, and degradation of long-chain fatty acids. Suppression of these transcripts correlates with increased intracellular lipid accumulation and reduced lipid catabolism identified by imaging and biochemical assays. The data reinforce the metabolic phenotype of DGUOK deficiency, linking nucleotide imbalance to impaired lipid utilization.

| Gene Symbol | Protein Encoded | Rank in Gene List | Rank Metric Score | Running Enrichment Score | Core Enrichment |
|---|---|---|---|---|---|

| | | | | | |
|---|---|---|---|---|---|
| *MT2A* | Metallothionein-2 A (metal/oxidative stress binder) | 3 | 124.6383 | 0.0013 | Yes |
| *CXCL8* | Interleukin-8 / CXCL8 chemokine (neutrophil chemoattractant) | 4 | 109.3565 | 0.0026 | Yes |
| *IL11* | Interleukin-11 cytokine | 11 | 76.6990 | 0.0038 | Yes |
| *JUN* | c-Jun, AP-1 transcription factor | 21 | 50.0915 | 0.0049 | Yes |
| *SHC1* | SHC adaptor protein 1 (RTK/STAT signaling adaptor) | 32 | 40.5686 | 0.0060 | Yes |
| *GRB10* | GRB10 adaptor (modulates RTK/insulin signaling) | 40 | 33.0000 | 0.0072 | Yes |
| *TIMP1* | Tissue inhibitor of metalloproteinases -1 | 41 | 31.5086 | 0.0085 | Yes |

| | | | | | |
|---|---|---|---|---|---|
| *CCND1* | Cyclin D1 (G1/S cell-cycle regulator) | 44 | 29.6576 | 0.0098 | Yes |
| *DUSP4* | Dual-specificity phosphatase 4 (MAPK phosphatase) | 54 | 25.5229 | 0.0109 | Yes |
| *PSMD2* | 26S proteasome regulatory subunit, non-ATPase 2 | 57 | 25.0555 | 0.0122 | Yes |
| *DUSP6* | MAPK phosphatase 6 (ERK-specific) | 68 | 21.2518 | 0.0133 | Yes |
| *FYN* | Src-family tyrosine kinase Fyn | 75 | 19.8539 | 0.0145 | Yes |
| *SQSTM1* | Sequestosome-1/p 62 (autophagy & NF-κB adaptor) | 87 | 18.6198 | 0.0155 | Yes |
| *EGR1* | Early growth response 1 transcription factor | 91 | 18.1427 | 0.0168 | Yes |
| *CNN2* | Calponin-2 (actin-binding) | 94 | 17.8539 | 0.0181 | Yes |

| | | | | | |
|---|---|---|---|---|---|
| *OAS1* | 2′-5′-oligoadenylat e synthetase 1 (antiviral) | 148 | 13.8861 | 0.0179 | Yes |
| *TUBA1A* | Tubulin alpha-1A | 157 | 13.1024 | 0.0190 | Yes |
| *JUNB* | JunB, AP-1 transcription factor | 172 | 12.4437 | 0.0200 | Yes |
| *BCL2L1* | Bcl-xL (anti-apoptotic mitochondrial protein) | 175 | 12.3372 | 0.0213 | Yes |
| *CFL1* | Cofilin-1 (actin depolymerizing factor) | 227 | 10.2924 | 0.0211 | Yes |

**Supplementary Table 3-H. Reactome "Cytokine Signaling in Immune System" — Core Enrichment (Top 20).**

This table summarizes the 20 most positively ranked genes driving enrichment of the cytokine signaling pathway. Genes such as *IL11*, *CXCL8*, *JUN*, *EGR1*, *OAS1*, and *TUBA1A* are among the highest contributors, collectively reflecting activation of STAT/MAPK-dependent transcription, chemokine production, and cytoskeletal reorganization. The coordinated upregulation of *GRB10*, *FYN*, and *SHC1* underscores enhanced receptor-tyrosine kinase and cytokine cross-talk, consistent with IFN-β–driven transcriptional reprogramming observed upon DGUOK knockdown. These data demonstrate that purine-driven interferon signaling extends to secondary cytokine cascades influencing cell growth and survival.

| Gene Symbol | Rank in Gene List | Rank Metric Score | Running Enrichment Score | Core Enrichment |
|---|---|---|---|---|
| *IDI2* | 14903 | 0.00000000 | -0.40359333 | No |
| *GGPS1* | 26866 | -0.28399664 | -0.72023140 | No |
| *PLPP6* | 29259 | -0.58502668 | -0.75391760 | No |
| *LBR* | 31178 | -1.31875873 | -0.77358920 | Yes |
| *PMVK* | 31234 | -1.35654736 | -0.73817830 | Yes |
| *ARV1* | 31829 | -1.82390869 | -0.71870380 | Yes |
| *SREBF1* | 31866 | -1.88605666 | -0.68273115 | Yes |
| *IDI1* | 32579 | -3.00000000 | -0.66674550 | Yes |
| *HSD17B7* | 32804 | -3.65757728 | -0.63633140 | Yes |
| *EBP* | 32847 | -3.82390881 | -0.60053610 | Yes |
| *SREBF2* | 32850 | -3.82390881 | -0.56355820 | Yes |
| *TM7SF2* | 32985 | -4.42021656 | -0.53048310 | Yes |

| | | | | |
|---|---|---|---|---|
| *NSDHL* | 33004 | -4.53760195 | -0.49397826 | Yes |
| *SC5D* | 33089 | -5.04575729 | -0.45942482 | Yes |
| *CYP51A1* | 33234 | -6.08618593 | -0.42664537 | Yes |
| *MVK* | 33250 | -6.23657179 | -0.39005184 | Yes |
| *FDFT1* | 33515 | -10.03151703 | -0.36082035 | Yes |
| *FDPS* | 33606 | -13.00877380 | -0.32644433 | Yes |
| *SQLE* | 33678 | -17.25963783 | -0.29150650 | Yes |
| *DHCR7* | 33688 | -18.30102921 | -0.25473556 | Yes |
| *MSMO1* | 33692 | -19.00000000 | -0.21778722 | Yes |
| *DHCR24* | 33710 | -20.22914886 | -0.18125282 | Yes |
| *ACAT2* | 33721 | -21.30102921 | -0.14451145 | Yes |
| *MVD* | 33722 | -21.35654640 | -0.10747441 | Yes |
| *LSS* | 33751 | -25.22184944 | -0.07126524 | Yes |

| | | | | |
|---|---|---|---|---|
| *HMGCR* | 33807 | -36.82390976 | -0.03585436 | Yes |
| *HMGCS1* | 33846 | -147.55284119 | 0.00005915 | Yes |

**Supplementary Table 4-I. Reactome "Cholesterol Biosynthesis" Core Enrichment (Top 20)**

This table lists the top negatively enriched genes within the Cholesterol Biosynthesis pathway, including SREBF1, PMVK, IDI2, GGPS1, LBR, and ARV1. These enzymes and regulators comprise the mevalonate pathway and sterol-assembly machinery essential for hepatic lipid balance. Their coordinated repression corresponds with decreased sterol and isoprenoid synthesis observed in transcriptomic data, reinforcing that DGUOK deficiency dampens anabolic lipid biosynthesis concurrently with accumulation of neutral lipids and interferon activation.

| Gene Symbol | Protein Encoded | Rank in Gene List | Rank Metric Score | Running Enrichment Score | Core Enrichment |
|---|---|---|---|---|---|
| *OAS1* | 2′–5′-Oligoadenylate synthetase 1 | 148 | 13.88605690 | 0.0023651960 | Yes |
| *TUBA1A* | Tubulin alpha-1A | 157 | 13.10237312 | 0.0088845715 | Yes |
| *TUBB2B* | Tubulin beta-2B | 255 | 9.72124672 | 0.0127630750 | Yes |
| *TUBB3* | Tubulin beta-3 | 269 | 9.43179798 | 0.0191340870 | Yes |
| *NUP58* | Nucleoporin 58 | 311 | 8.46852112 | 0.0246742630 | Yes |
| *UBC* | Polyubiquitin-C | 322 | 8.34678745 | 0.0311342920 | Yes |
| *PLCG1* | Phospholipase C gamma-1 | 435 | 6.46852112 | 0.0345677060 | Yes |
| *TUBB6* | Tubulin beta-6 | 444 | 6.39794016 | 0.0410870800 | Yes |
| *PPP2R5A* | PP2A regulatory subunit B′ alpha | 452 | 6.31875896 | 0.0476361300 | Yes |
| *TUBA1C* | Tubulin alpha-1C | 469 | 6.17392540 | 0.0539181230 | Yes |
| *NUP43* | Nucleoporin 43 | 519 | 5.69896984 | 0.0592209180 | Yes |

| | | | | | |
|---|---|---|---|---|---|
| *HSPA2* | Heat shock protein 70-2 | 707 | 4.30980396 | 0.0604288760 | Yes |
| *TUBA1B* | Tubulin alpha-1B | 772 | 4.09151506 | 0.0652865800 | Yes |
| *UBE2I* | UBC9 (SUMO-conjugating enzyme E2 I) | 905 | 3.61978865 | 0.0681265400 | Yes |
| *PPP2CA* | PP2A catalytic subunit alpha | 930 | 3.55284190 | 0.0741711500 | Yes |
| *KPNA4* | Importin alpha-3 | 1026 | 3.25963736 | 0.0781090000 | Yes |
| *CENPX* | Centromere protein X | 1065 | 3.16749120 | 0.0837381900 | Yes |
| *ISG15* | ISG15 ubiquitin-like modifier | 1076 | 3.14266753 | 0.0901982260 | Yes |
| *FLNA* | Filamin-A | 1252 | 2.74472761 | 0.0917622600 | Yes |
| *UBE2E1* | Ubiquitin-conjugating enzyme E2 E1 | 1341 | 2.58502674 | 0.0959078150 | Yes |

**Supplementary Table 4-I. Reactome "Fatty Acid β-Oxidation and Peroxisomal Transport"**

This pathway specifically resolves the subset of genes within long-chain β-oxidation and peroxisomal transport processes, separating it from global lipid metabolism. Down-regulated core genes such as SLC27A3, SLC25A17, HACD2, MCAT, and MLYCD indicate suppression of fatty-acid activation, elongation, and CoA transport machinery. Reduced expression of PCTP and HPGD suggests disrupted phospholipid and prostaglandin handling. Collectively, these changes point to impaired peroxisome–mitochondria coordination in energy and lipid homeostasis following DGUOK loss.

| Gene Symbol | Protein Encoded | Rank in Gene List | Rank Metric Score | Running Enrichment Score | Core Enrichment |
|---|---|---|---|---|---|
| *POLR1D* | RNA polymerase I and III subunit D (transcription machinery) | 243 | 10.036 | –0.0019 | Yes |
| *SMARCD1* | SWI/SNF chromatin remodeling complex subunit 1 | 359 | 7.538 | –0.00005 | Yes |
| *DR1* | Down-regulator of transcription 1 (NC2β repressor) | 601 | 5.000 | –0.0019 | Yes |
| *UBE2I* | SUMO-conjugating enzyme E2 I (UBC9) | 905 | 3.620 | –0.0056 | Yes |
| *POLR2K* | RNA polymerase II subunit K | 931 | 3.553 | –0.0011 | Yes |
| *SMARCC1* | SWI/SNF chromatin remodeling complex subunit C1 | 1064 | 3.170 | –0.0021 | Yes |
| *TAF10* | TFIID subunit TAF10 (transcription initiation factor) | 1072 | 3.153 | –0.0022 | Yes |

| | | | | | |
|---|---|---|---|---|---|
| *HAT1* | Histone acetyltransferase type B catalytic subunit 1 | 1088 | 3.115 | –0.0024 | Yes |
| *NUP107* | Nucleoporin 107 (karyopherin complex scaffold) | 1154 | 2.948 | –0.0025 | Yes |
| *SUPT5H* | Transcription elongation factor Spt5 | 1214 | 2.803 | –0.0026 | Yes |
| *RUVBL2* | AAA+ ATPase component of chromatin remodeling complexes | 1232 | 2.764 | –0.0027 | Yes |
| *H2AFZ* | Histone variant H2A.Z (promotes nucleosome mobility) | 1242 | 2.742 | –0.0028 | Yes |
| *SUPT16H* | FACT complex subunit Spt16 (histone chaperone) | 1285 | 2.656 | –0.0029 | Yes |
| *TAF9* | TFIID subunit TAF9 (transcription initiation factor) | 1310 | 2.608 | –0.0030 | Yes |
| *RUVBL1* | AAA+ ATPase RuvB-like 1 (chromatin remodeler) | 1321 | 2.586 | –0.0031 | Yes |

| | | | | | |
|---|---|---|---|---|---|
| *CHD1* | Chromodomain-helicase-DNA-binding protein 1 (remodeler) | 1333 | 2.562 | –0.0031 | Yes |
| *TAF5* | TFIID subunit TAF5 (core scaffold component) | 1339 | 2.550 | –0.0032 | Yes |
| *ING3* | Inhibitor of growth family member 3 (NuA4 complex subunit) | 1343 | 2.542 | –0.0033 | Yes |
| *RBBP7* | Histone-binding protein RbAp46/48 (chromatin assembly factor) | 1349 | 2.530 | –0.0033 | Yes |
| *EP400* | E1A-binding protein p400 (Tip60 HAT complex component) | 1352 | 2.524 | –0.0034 | Yes |

**Supplementary Table 4-J. Reactome "Epigenetic Regulation of Gene Expression"**

This table lists the top 20 leading-edge genes contributing to enrichment of the Reactome *"Epigenetic Regulation of Gene Expression"* pathway identified by GSEA preranked analysis comparing **siDGUOK** and **siCON HEPG2** cells. Core genes such as *SMARCD1*, *RUVBL1*, *RUVBL2*, *EP400*, and *CHD1* encode components of chromatin-remodeling and transcriptional coactivator complexes including SWI/SNF, NuA4, and TFIID. The listed genes exhibit positive rank-metric scores, reflecting relative upregulation in the siDGUOK condition. Enrichment within this pathway indicates increased representation of factors involved in histone modification, nucleosome remodeling, and transcriptional regulation under the analyzed condition.

| Gene Symbol | Rank in Gene List | Rank Metric Score | Running Enrichment Score | Core Enrichment |
|---|---|---|---|---|
| *POLR1D* | 243 | 10.0362 | −0.0019 | Yes |
| *SMARCD1* | 359 | 7.5376 | −0.00005 | Yes |
| *DR1* | 601 | 5.0000 | −0.0019 | Yes |
| *UBE2I* | 905 | 3.6198 | −0.0056 | Yes |
| *POLR2K* | 931 | 3.5528 | −0.0011 | Yes |
| *SMARCA4* | 1078 | 3.1367 | −0.0001 | Yes |
| *RBM7* | 1103 | 3.0605 | 0.0044 | Yes |
| *DEK* | 1129 | 3.0000 | 0.0090 | Yes |
| *ATF7IP* | 1143 | 2.9586 | 0.0139 | Yes |
| *SPOCD1* | 1347 | 2.5686 | 0.0132 | Yes |
| *SUMO2* | 1364 | 2.5376 | 0.0180 | Yes |
| *ZNF765* | 1607 | 2.1938 | 0.0161 | Yes |
| *ZNF669* | 1631 | 2.1739 | 0.0207 | Yes |
| *ACTB* | 1749 | 2.0605 | 0.0225 | Yes |

| | | | | |
|---|---|---|---|---|
| *POLR2H* | 1826 | 2.0000 | 0.0255 | Yes |
| *ZNF264* | 2269 | 1.6021 | 0.0177 | Yes |
| *SS18L1* | 2424 | 1.5229 | 0.0184 | Yes |
| *SAP30* | 2453 | 1.5086 | 0.0229 | Yes |
| *SETD1B* | 2542 | 1.4559 | 0.0255 | Yes |
| *MTA2* | 2606 | 1.4202 | 0.0290 | Yes |

**Supplementary Table 4-J. Reactome "Epigenetic Regulation of Gene Expression"**

This table lists the top 20 leading-edge genes contributing to enrichment of the Reactome "Epigenetic Regulation of Gene Expression" pathway identified by GSEA preranked analysis comparing siDGUOK and siCON HEPG2 cells. Core enriched genes include POLR1D, SMARCD1, SMARCA4, UBE2I, and SETD1B, which encode components of chromatin-remodeling, histone-modifying, and transcriptional co-regulatory complexes. The listed genes exhibit positive rank-metric scores, indicating relative upregulation in the siDGUOK condition. Enrichment within this pathway denotes increased representation of genes associated with histone modification, nucleosome remodeling, and transcriptional regulation in the analyzed comparison.

| Gene Symbol | Rank in Gene List | Rank Metric Score | Running Enrichment Score | Core Enrichment |
|---|---|---|---|---|
| *MRI1* | 1137 | 2.9586 | 0.1331 | Yes |
| *MTAP* | 1478 | 2.3566 | 0.2897 | Yes |
| *ENOPH1* | 2041 | 1.7696 | 0.4398 | Yes |
| *GOT1* | 2942 | 1.2366 | 0.5798 | Yes |
| *APIP* | 8122 | 0.3010 | 0.5935 | Yes |
| *ADI1* | 33585 | −12.3565 | 0.0078 | No |

**Supplementary Table 4-K. Reactome "Methionine Salvage Pathway"**

This table lists the leading-edge genes contributing to enrichment of the Reactome *"Methionine Salvage Pathway"* identified by GSEA preranked analysis comparing siDGUOK and siCON HEPG2 cells. Core enriched genes such as *MRI1*, *MTAP*, *ENOPH1*, *GOT1*, and *APIP* encode enzymes involved in the methionine salvage and transsulfuration cycles. The listed genes exhibit positive rank-metric scores, indicating relative upregulation in the siDGUOK condition. Enrichment within this pathway reflects increased representation of genes involved in methionine recycling, polyamine metabolism, and related sulfur-amino-acid processes under the analyzed comparison.

| Gene Symbol | Rank in Gene List | Rank Metric Score | Running Enrichment Score | Core Enrichment |
|---|---|---|---|---|
| *PNP* | 84 | 18.7212 | 0.0083 | No |
| *GLRX* | 133 | 14.4559 | 0.0176 | No |
| *GUK1* | 289 | 8.9208 | 0.0238 | No |
| *CDA* | 400 | 6.7959 | 0.0312 | No |
| *GART* | 748 | 4.1739 | 0.0317 | No |
| *AK6* | 874 | 3.6990 | 0.0388 | No |
| *NME2* | 897 | 3.6576 | 0.0489 | No |
| *DCTPP1* | 1043 | 3.2366 | 0.0553 | No |
| *TXNRD1* | 1107 | 3.0555 | 0.0642 | No |
| *NME3* | 1377 | 2.5086 | 0.0670 | No |
| *TXN* | 1496 | 2.3279 | 0.0743 | No |
| *RRM2* | 1569 | 2.2366 | 0.0829 | No |
| *DCK* | 1650 | 2.1487 | 0.0913 | No |
| *NME1* | 1688 | 2.1192 | 0.1009 | No |

| | | | | |
|---|---|---|---|---|
| *GMPR2* | 1703 | 2.0969 | 0.1113 | No |
| *ENTPD6* | 1893 | 1.9208 | 0.1164 | No |
| *NUDT15* | 1937 | 1.8539 | 0.1259 | No |
| *CTPS1* | 2048 | 1.7696 | 0.1334 | No |
| *AK2* | 2103 | 1.7213 | 0.1425 | No |
| *AMPD3* | 2169 | 1.6778 | 0.1514 | No |

**Supplementary Table 4-L. Reactome "Metabolism of Nucleotides"**

This table lists the top 20 ranked genes contributing to the Reactome *"Metabolism of Nucleotides"* pathway identified by GSEA preranked analysis comparing siDGUOK and siCON HEPG2 cells. Genes such as *PNP*, *GUK1*, *CDA*, *NME1*, and *CTPS1* encode enzymes involved in nucleotide interconversion, deoxyribonucleotide synthesis, and purine–pyrimidine metabolism. The rank-metric scores are positive, indicating relative upregulation in the siDGUOK condition, while none of the listed genes were designated as core enriched within the leading-edge subset. This enrichment profile represents the relative contribution of nucleotide metabolic genes to the ranked gene list within the analyzed comparison.

| Gene Symbol | Rank in Gene List | Rank Metric Score | Running Enrichment Score | Core Enrichment |
|---|---|---|---|---|
| *PNP* | 84 | 18.7212 | 0.0269 | No |
| *ENTPD6* | 1893 | 1.9208 | 0.0029 | No |
| *NUDT15* | 1937 | 1.8539 | 0.0310 | No |
| *ITPA* | 3021 | 1.2076 | 0.0284 | No |
| *ADPRM* | 3326 | 1.0809 | 0.0488 | No |
| *NT5C* | 4301 | 0.7959 | 0.0494 | No |
| *UPP1* | 4343 | 0.7959 | 0.0776 | No |
| *ENTPD4* | 4563 | 0.7447 | 0.1006 | No |
| *NUDT18* | 5746 | 0.5376 | 0.0950 | No |
| *DPYS* | 9967 | 0.1805 | −0.0004 | No |
| *UPP2* | 10898 | 0.1308 | 0.0015 | No |
| *NT5C3A* | 11717 | 0.1079 | 0.0068 | No |
| *ENTPD1* | 11954 | 0.0915 | 0.0292 | No |
| *TYMP* | 12159 | 0.0757 | 0.0526 | No |

| | | | | |
|---|---|---|---|---|
| *ENTPD3* | 12696 | 0.0506 | 0.0661 | No |
| *XDH* | 15031 | 0.0000 | 0.0265 | No |
| *NT5C1B* | 15754 | 0.0000 | 0.0346 | No |
| *NUDT5* | 25301 | −0.1308 | −0.2183 | No |
| *NT5M* | 25819 | −0.1739 | −0.2042 | No |
| *AGXT2* | 27221 | −0.3188 | −0.2162 | No |

**Supplementary Table 4-M. Reactome "Nucleotide Catabolism"**

This table lists the top 20 ranked genes contributing to the Reactome *"Nucleotide Catabolism"* pathway identified by GSEA preranked analysis comparing **siDGUOK** and **siCON HEPG2** cells. Genes such as *PNP*, *ITPA*, *ADPRM*, *ENTPD* family members, and *NT5C* encode enzymes involved in nucleotide dephosphorylation, base salvage, and degradation. All listed genes exhibit a range of positive to negative rank-metric scores, indicating variable relative expression across the ranked gene list. None of the genes in this set were designated as core enriched, and the table reflects the distribution of catabolic enzymes contributing to overall pathway representation.

| Gene Symbol | Rank in Gene List | Rank Metric Score | Running Enrichment Score | Core Enrichment |
|---|---|---|---|---|
| *STAT4* | 148 | 15.231 | 0.0112 | Yes |
| *IRF1* | 297 | 9.875 | 0.0283 | Yes |
| *JAK2* | 352 | 8.472 | 0.0347 | Yes |
| *SOCS3* | 488 | 6.312 | 0.0465 | Yes |
| *IFNG* | 703 | 4.219 | 0.0518 | Yes |
| *BCL3* | 745 | 3.998 | 0.0601 | Yes |
| *IL12RB1* | 912 | 3.515 | 0.0682 | Yes |
| *IL12RB2* | 954 | 3.421 | 0.0719 | Yes |
| *TYK2* | 1068 | 3.104 | 0.0807 | Yes |
| *IFNGR1* | 1101 | 3.018 | 0.0833 | Yes |
| *NFKBIA* | 1143 | 2.941 | 0.0912 | Yes |
| *CISH* | 1267 | 2.714 | 0.0989 | Yes |

| | | | | |
|---|---|---|---|---|
| *JUNB* | 1362 | 2.552 | 0.1034 | Yes |
| *IL27RA* | 1430 | 2.413 | 0.1117 | Yes |
| *TNFAIP3* | 1478 | 2.358 | 0.1202 | Yes |
| *IRF8* | 1555 | 2.236 | 0.1265 | Yes |
| *BATF* | 1628 | 2.146 | 0.1336 | Yes |
| *NMI* | 1730 | 2.043 | 0.1397 | Yes |
| *SOCS1* | 1824 | 1.956 | 0.1473 | Yes |
| *FOS* | 1902 | 1.889 | 0.1556 | Yes |

**Supplementary Table 4-N. Reactome "Gene and Protein Expression by JAK STAT Signaling After Interleukin 12 Stimulation"**

This table lists the top 20 leading-edge genes contributing to enrichment of the Reactome *"Gene and Protein Expression by JAK-STAT Signaling after Interleukin-12 Stimulation"* pathway identified by GSEA preranked analysis comparing **siDGUOK** and **siCON HEPG2** cells. Core enriched genes include *STAT4*, *IRF1*, *JAK2*, *SOCS1*, and *IFNG*, representing canonical members of the IL-12–JAK–STAT regulatory axis. The listed genes exhibit positive rank-metric scores, indicating relative upregulation in the siDGUOK condition. Enrichment within this pathway reflects increased representation of transcripts associated with cytokine-dependent JAK–STAT signal transduction and transcriptional responses to interleukin stimulation in the analyzed comparison.
